# Self-generated action shapes theta phase alignment at encoding and later recollection

**DOI:** 10.64898/2026.11.18.767418

**Authors:** Qin Chen, Jie Sui

## Abstract

Across species, self-directed action is linked to theta rhythms and better memory, but such action combines choosing, moving, and controlling the outcome. Whether choosing and control over the outcome shape recollection, and the encoding dynamics that support it, remains unclear. Three preregistered experiments in human participants separated these components while holding movement constant. Participants encoded scene pairs by dragging either a freely chosen scene or one indicated by the program, with fluent or disrupted feedback. Recognition was tested after 24 hours (and immediately in Experiment 2), using Remember/Know judgments. Experiment 3 recorded scalp electroencephalography during encoding. Choosing improved recognition, more for recollection than familiarity. A smaller benefit extended to the unselected scene, although less consistently after latency adjustment. Disrupted feedback lowered rated agency but did not reduce memory at 24 hours. In Experiment 2, however, recollection declined less over 24 hours after self-generated than externally generated action, but only under fluent feedback. Choosing increased action-onset-locked theta and alpha phase alignment (inter-trial phase coherence), whereas disrupted feedback reduced alignment later. The selection effects survived matching for response-initiation-latency. Trial by trial, later posterior theta alignment (353–570 ms) predicted Remember over Know responses, whereas early frontal alpha alignment (127–245 ms) predicted Remember over Forgotten responses. These findings link self-generated action to the timing of encoding-related oscillations and to subsequent recollection. The association between later posterior theta alignment and Remember–Know discrimination is consistent with a role for theta phase coordination in associative binding.

**Significance Statement:** People often remember events they bring about better than events they only observe. Acting, however, involves several steps: choosing what to do, doing it, and seeing its result. Three experiments separated these steps while keeping movement identical. Choosing which scene to act on improved memory for it, most for recollection, the detailed re-experiencing of an event. Smooth control over the outcome did not create this benefit but helped it last over a day. In brain recordings, choosing and outcome feedback changed the alignment of brain rhythms to the action at different moments. Later alignment distinguished recollected events from those that only felt familiar. Part of what makes a memory one’s own may therefore be set at the moment of choosing.

## Introduction

Recollecting an episode means re-experiencing it as one’s own past, not merely knowing that it occurred. Tulving termed this first-person quality autonoetic consciousness (Tulving, 1985, 2002). Whether it is shaped only at retrieval or partly at encoding, by being the agent of an action, remains unclear. Self-directed behavior links action to memory, but choosing, moving, and controlling an outcome are usually intertwined.

Across species, self-directed behavior is linked to theta rhythms and better memory. In rats, theta rhythms accompany voluntary movement (Vanderwolf, 1969), and place-cell firing depends on self-movement (Lu and Bilkey, 2010). In monkeys, saccades during visual exploration reset theta phase, and more reliable resetting is associated with better subsequent recognition (Jutras et al., 2013). In humans, volitional exploration improves memory in healthy adults but not amnesic patients (Voss et al., 2011). Choosing also improves recognition when it does not change what is seen (Murty et al., 2015; DuBrow et al., 2019). These findings suggest a link between self-directed action, oscillatory timing, and memory, but leave unclear how different components of agency shape encoding dynamics relevant to recollection.

The sense of agency is the experience of causing and controlling one’s own actions and their consequences (Haggard, 2017). Accounts distinguish prospective signals arising during action selection from retrospective signals arising from the match between predicted and actual sensory outcomes (Wenke et al., 2010; Haggard and Chambon, 2012; Sidarus et al., 2013; Chambon et al., 2014). Because the two signals arise at different stages of an action, they can be varied independently.

Whether these memory benefits reflect recollection or familiarity also remains unclear. In dual-process accounts, recollection depends on binding episodic elements, whereas familiarity reflects graded memory strength (Yonelinas, 2002). Theta phase is proposed to organize associative binding (Nyhus and Curran, 2010; Backus et al., 2016; Clouter et al., 2017). The theta–gamma neural-code account proposes that gamma-associated representations occupy different phases of a theta cycle (Lisman and Jensen, 2013). Theta–gamma coupling increases during item-context learning in rats and is associated with temporal-order memory in humans (Tort et al., 2009; Heusser et al., 2016). Phase alignment and coupling therefore offer complementary measures of encoding dynamics as an action unfolds.

Three questions follow. First, which part of agency (choosing, performing, or controlling the outcome) affects memory, which kind of memory benefits, and does the benefit extend beyond the chosen item? A benefit of performing an action should not differ between chosen and assigned actions when movement is matched. A benefit of choosing should follow selection. If the experience of control marks actions as one’s own in memory (Wegner, 2004), the benefit should track rated agency and weaken with disrupted feedback. A larger recollection benefit would be consistent with stronger episodic binding. Extension to the unselected item would suggest an effect beyond the chosen object. Second, does the benefit change between an immediate and a 24-hour test? Third, are different components of agency expressed in temporally distinct neural dynamics at encoding, and do these dynamics predict later memory? If choosing strengthens binding, theta phase alignment at encoding should track later recollection selectively. Theta–gamma phase–amplitude coupling (PAC) offers a further test of the binding account. If alpha phase alignment instead reflects attention to the items, it should track recognition more broadly.

Three preregistered experiments crossed action selection with action feedback while matching movement. Participants dragged either their chosen scene (self-generated) or the program-indicated scene (externally generated), with fluent or disrupted feedback. These manipulations targeted prospective and retrospective agency signals, respectively (Sato and Yasuda, 2005; Wenke et al., 2010). Explicit agency ratings and time-compression estimates assessed their subjective effects (Synofzik et al., 2008; Schwarz et al., 2019; Lafleur et al., 2020). Remember/Know judgments distinguished reported recollection from familiarity. Unselected scenes were also tested. All experiments tested memory after 24 hours. Experiment 2 added an immediate test. Experiment 3 recorded scalp electroencephalography (EEG) to examine the timing, frequency, and trial-wise memory relevance of encoding dynamics.

## Materials and Methods

### Ethics statement

All procedures were approved by the University of Aberdeen ethics committee (ID: 3320512). Written informed consent was obtained from all participants prior to testing.

### Participants

Participants were recruited from the University of Aberdeen School of Psychology participant pool (SONA, and ORSEE for Experiment 3) and received course credit or monetary compensation. Inclusion criteria were age 18–45 years and normal or corrected-to-normal vision. For Experiment 3, participants were additionally excluded for self-reported neurological illness or prior head injury involving loss of consciousness.

A pilot behavioral sample (n = 13; mean age = 20.31 ± 2.78 years; 12 women) and an EEG feasibility sample (n = 8; mean age = 30.63 ± 10.77 years; 5 women) were collected to optimize task parameters and estimate effect sizes. Pilot and feasibility data were not included in the confirmatory analyses reported here. Recruitment for the three experiments followed the fixed-sample-size targets specified in the preregistration.

In Experiment 1, 47 participants were tested and 1 was excluded for chance-level memory performance, leaving 46 participants (mean age = 20.41 ± 2.42 years; 40 women, 6 men). In Experiment 2, all 47 participants were tested, and all met the prespecified overall recognition-performance criterion, leaving 47 participants for analysis (mean age = 20.38 ± 3.65 years; 35 women, 11 men, 1 non-binary). In Experiment 3, 46 participants were tested, of whom 3 were excluded for chance-level memory performance and 1 for missing EEG recording, leaving 42 participants (mean age = 22.21 ± 3.41 years; 31 women, 11 men) for the final EEG analyses. For two participants with minor recording loss, all available valid trials from these participants were retained in the EEG analyses. Retained action-onset-locked EEG trial counts by experimental condition and recognition response are summarized in Tables S1 and S2 in Text S1, and participant-level counts are provided on OSF (https://osf.io/3yhpz).

### Stimuli and apparatus

Stimuli were color pictures depicting indoor and outdoor scenes. Images were drawn primarily from the SUN database and supplemented with freely available high-quality pictures from online repositories. A total pool of 490 images, including practice items, was used to build the encoding and recognition lists. The tasks were run in PsychoPy (version 22.2.3). Displays had a resolution of 1920 × 1080 pixels, and viewing distance was 80 cm in all three experiments. Scenes subtended approximately 7.24° × 7.24° of visual angle across experiments.

### Experimental design and statistical analysis

Experiments 1 and 3 used a within-participant 2 × 2 factorial design at encoding, crossing action selection (self-generated vs. externally generated) with action feedback (fluent vs. disrupted). In the self-generated condition, participants freely chose which of two simultaneously presented images to move, whereas in the externally generated condition, the target image was indicated by the program. Action feedback was manipulated by varying the image-following coefficient during dragging (.09 for fluent trials and .01 for disrupted trials). At each update, the image moved 9% or 1% of the remaining distance to the current mouse position, respectively. Fluent trials therefore had a shorter movement-feedback lag and higher controllability than disrupted trials. Experiment 2 used the same encoding manipulations and tested memory immediately and after a 24 h delay.

Primary behavioral outcomes were analyzed using Bayesian mixed-effects models. Condition-level EEG effects were evaluated using cluster-based permutation tests, followed by trial-wise neural–memory modeling, exploratory decoding, and focused moderation analyses. Detailed procedures are provided below under “Behavioral measures and statistical analyses,” “EEG feature extraction and inferential analyses,” and “Trial-wise prediction analyses (Experiment 3).”

### Procedure

#### Encoding task

On each encoding trial, two scene images were presented simultaneously on the left and right sides of the screen. A cue indicated whether action selection was self-generated or externally generated. In self-generated trials, participants freely selected one image to move, whereas in externally generated trials, the program highlighted the image to move. Participants moved the selected image using a continuous dragging action until it disappeared from the screen. During movement, the action feedback manipulation produced fluent or disrupted visual feedback, with smaller or larger movement-feedback lags, respectively. Following each drag, participants first estimated the duration of scene movement, from which the time-compression measure was derived as an indirect index of agency. They then rated their explicit sense of agency on a 0–100 scale. The encoding task comprised four blocks and a total of 164 trials (328 images; two images per trial), including four practice trials. Trial order was pseudorandomized within blocks, with the constraint that no more than three consecutive trials belonged to the same Selection × Feedback condition. Participants were asked to perform the task accurately and were not told about the later memory test.

#### Remember/Know/New test

Recognition memory was assessed with a standard Remember/Know/New procedure (Elliott et al., 2020). On each recognition trial, a single image was presented centrally, and participants judged whether the image was new, familiar in the absence of specific contextual details (Know), or remembered together with specific details from the encoding episode (Remember). Participants selected one of five judgments: Definitely New, Maybe New, Maybe Know, Definitely Know, or Remember. For categorical recognition analyses, the two New judgments were combined, as were the two Know judgments. Remember responses formed a separate category. In Experiment 1, the recognition test was conducted approximately 24 h after encoding and comprised 244 trials (4 practice trials, then 160 old and 80 new images). Experiment 2 included both an immediate recognition test and a delayed test conducted approximately 24 h later with 244 trials per test (four practice trials, then 160 old and 80 new images). The immediate and delayed tests used non-overlapping sets of encoded scenes. Experiment 3 used the same recognition procedure as Experiment 1 but included a larger test set of 364 trials in total (4 practice trials, followed by 240 old and 120 new images), divided into 6 sessions. Across all experiments, trial order was pseudorandomized within each recognition-test session. No more than three consecutive trials belonged to the same experimental condition or stimulus category.

#### Unselected-item recognition

Unselected scenes were also tested at recognition to assess whether the effects of the encoding manipulations extended beyond selected items. These trials were analyzed separately.

### Behavioral measures and statistical analyses

Primary outcomes followed the preregistered unit of analysis. For old items, New responses were coded as Forgotten in the recognition models. In Experiments 1 and 3, trial-level recognition outcomes were analyzed using Bayesian multinomial mixed-effects models, with Know and Remember responses modeled relative to Forgotten responses. Unselected-item recognition was analyzed separately as correct versus incorrect old-item recognition. Selection, Feedback, and their interaction were entered as fixed effects.

In Experiment 2, corresponding multinomial and Bernoulli models included Selection, Feedback, Delay, and their interactions to analyze selected-item memory and unselected-item recognition, respectively.

In Experiment 2, immediate and delayed recognition were additionally used to derive retention ratios for recollection, familiarity and unselected-item recognition. Retention was defined as delayed performance divided by immediate performance within participant and encoding condition. Retention ratios were analyzed using Bayesian Student-*t* mixed-effects models with Selection, Feedback, and their interaction as predictors. A joint model additionally tested whether the Selection × Feedback effect differed between recollection- and familiarity-based retention.

To check that the manipulations changed agency, explicit and indirect indices of agency were analyzed. Explicit agency was indexed by sense-of-agency ratings standardized within participant. Time compression was calculated on each trial as the estimated image-movement duration minus the recorded movement duration, both in milliseconds. Negative values indicated underestimation and positive values indicated overestimation. Scores were z-standardized within participant before model fitting and used as an indirect index of agency. Both measures were analyzed using Bayesian mixed-effects models with fixed effects of Selection, Feedback, and their interaction.

Across Experiments 1–3, recognition discriminability was quantified as *d′*. Remember and Know *d′* were calculated from selected old items, whereas unselected-item *d′* was calculated from the corresponding unselected old items. False-alarm rates were estimated from new trials at the participant level, and rates were log-linear corrected before conversion to z-scores. Discriminability estimates were analyzed using Bayesian Student-t mixed-effects models with Selection, Feedback, and their interaction as fixed effects. Experiment 2 models also included Delay.

Know discriminability was additionally corrected using the independent Remember–Know (IRK) procedure, P(Know)/[1-P(Remember)] (Yonelinas and Jacoby, 1995; Song et al., 2011), before conversion to *d′*. A harmonized IRK-corrected Know analysis was also run across experiments.

For selected-item confidence data, equal-variance signal-detection (EVSD), unequal-variance signal-detection (UVSD), and dual-process signal-detection (DPSD) models were compared across the delayed tests of Experiments 1–3. The primary analysis included pooled data and experiment-specific fits. Models were compared using likelihood-based information criteria, leave-one-out predictive criteria, response-category goodness of fit, and posterior-predictive accuracy and coverage of the receiver operating characteristic (ROC).

Unless otherwise specified, Bayesian mixed-effects models were estimated in R using brms with four chains, 4,000 iterations per chain, and 2,000 warm-up iterations. Sampling used adapt_delta = .99 and max_treedepth = 14. Fixed-effect coefficients were assigned Normal (0, 1) priors, intercepts Student–*t* (3, 0, 2.5) priors, group-level standard deviations Exponential (1) priors, and random-effect correlation matrices LKJ (2) priors; Gaussian residual standard deviations received Student–*t* (3, 0, 2.5) priors. The Experiments 1–3 Student-t *d′* models instead used four chains with 6,000 iterations per chain and 3,000 warm-up iterations. Fixed effects received Normal (0, 0.5) priors, intercepts received Student-t (3, 0, 1) priors, group-level and residual standard deviations received Exponential (2) priors, and the degree-of-freedom parameters received Gamma (2, 0.1) priors. Prior predictive checks, posterior predictive checks, and two prespecified prior-sensitivity schemes were conducted for the confirmatory model set.

Convergence and sampling diagnostics were assessed using *R*, effective sample sizes, divergent transitions, maximum-treedepth events, and Pareto-k diagnostics where leave-one-out comparison was used. Bayesian estimates are reported as posterior means with 95% credible intervals (CrIs) in brackets.

#### Hierarchical drift-diffusion modeling (HDDM)

Hierarchical drift-diffusion models were fitted using HDDM v1.0.1RC to examine condition-dependent latent decision processes for selected-item Remember and Know responses and unselected-item old-new recognition. Selected-item models contrasted old items receiving the relevant target response (Remember or Know) with missed old items, whereas unselected-item models contrasted correctly recognized with missed unselected old items. For each experiment and outcome, eight candidate models (M1–M8) were fitted. All models allowed drift rate (*ν*) to vary by condition (Selection × Feedback in Experiments 1 and 3, and Selection × Feedback × Delay in Experiment 2), while additional models allowed non-decision time (*t*), starting point (*z*), boundary separation (*a*), or their combinations to vary by condition. All models included a 3% outlier-mixture probability (p_outlier = .03). Models were compared using the deviance information criterion (DIC), with lower values indicating better relative fit on the same observations. Four independent chains were run per model, each comprising 8,000 samples, with a burn-in of 1,200 and thinning of 2. Model M2 of the Experiment 2 unselected-item analysis used 12,000 samples and a burn-in of 2,000. Convergence was assessed using the Gelman–Rubin statistic (*R*), with the numbers of parameters exceeding 1.01 and 1.05 recorded for diagnostic inspection. Posterior contrasts were derived from the best-supported model for each experiment and outcome (Voss et al., 2013; White and Poldrack, 2014).

### EEG acquisition and preprocessing (Experiment 3)

EEG was recorded during the encoding phase of Experiment 3 using a 64-channel BioSemi ActiveTwo system with sintered Ag/AgCl electrodes at a sampling rate of 1024 Hz and stored for offline analysis. Preprocessing was performed in EEGLAB 2025.0.0 in MATLAB 2021b. Continuous data were first filtered using a zero-phase finite impulse response (FIR) filter (1–100 Hz) and a line-noise notch filter (49–51 Hz). Continuous recordings were then epoched from −0.5 to 2.0 s around action onset, the start of the selection-and-dragging movement. Baseline correction was applied using the −200 to 0 ms pre-action interval. Epoched data were re-referenced to the common average across channels. Independent component analysis (ICA, 45 components) was used to remove artifactual components (e.g., blinks/saccades and muscle activity), identified from their topographies and time courses. On average, about 9 components were rejected per participant (SD = 4.28; N = 42). No additional epoch-level trial-rejection procedure was applied after ICA. Each participant therefore contributed approximately 40 valid trials per Selection × Feedback condition to the EEG analyses, except for two participants with partial recording loss.

### EEG feature extraction and inferential analyses

#### Time-frequency decomposition and inter-trial phase coherence

Time-frequency representations were computed using complex Morlet wavelet convolution. ITPC was calculated from trial-wise phase angles to quantify the consistency of oscillatory phase across trials (Tallon-Baudry and Bertrand, 1999; Fell et al., 2008; Clouter et al., 2017; Lakatos et al., 2019; Guth et al., 2025; Long et al., 2025). At each frequency-time point, ITPC was defined as:

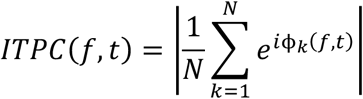

where *N* is the number of trials per condition per participant, and **φ***_k_*(*f*, *t*) = arg:*a_k_*(*f*, *t*); is the phase angle of the complex analytic signal *a_k_*(*f*, *t*) for trial *k* at frequency *f* and time *t*. Frequencies from 1 to 20 Hz were sampled on a logarithmic scale (38 frequencies; constant 4-cycle wavelets). For condition-level analyses, trials were grouped into the four conditions, corresponding to the 2 × 2 design (Selection × Feedback: self-generated–disrupted, self-generated–fluent, externally generated–disrupted, and externally generated–fluent).

Inferential analyses focused on the theta and alpha activity within the action-onset-locked 0–1 s interval. Frequency bins were selected nearest to the nominal theta (4–7.99 Hz) and alpha (8– 12.99 Hz) boundaries. The resulting bin ranges were approximately 3.96–8.21 Hz and 8.21–13.34 Hz, respectively, so the 8.21 Hz bin contributed to both bands. For each band, ITPC was averaged across frequencies within the band, and inference used two-dimensional non-parametric cluster-based permutation tests (CBPTs) across channels and time using FieldTrip toolbox (Oostenveld et al., 2011). Separate tests evaluated the main effects of Selection, Feedback, and their interaction. Cluster formation used a sample-level threshold of *p* < .001, with cluster significance assessed at *p* < .05 using 10,000 Monte Carlo randomizations. Spatial clustering was defined using distance-based channel neighborhoods, with a minimum of three neighboring channels required for cluster formation. FieldTrip cluster correction controlled the family-wise error rate across the whole channel × time search space within each frequency band. For each effect, the minimum cluster-level *p* values from the theta and alpha bands were additionally adjusted using the Benjamini–Hochberg false-discovery-rate (BH–FDR) procedure across the two prespecified bands. For concise reporting and visualization, the first-ranked FieldTrip cluster (cluster 1) was selected within each main effect and frequency band. Clusters shorter than 50 ms were not interpreted further. Cluster extents in time are reported descriptively and are not precise estimates of effect onset or offset. Regional summaries were used only to describe significant whole-head clusters and to construct the subsequent trial-wise predictors. All condition-level ITPC analyses and visualizations used raw ITPC values without baseline correction.

#### Trial-wise jackknife inter-trial phase-coherence features (jITPC)

To link encoding-related neural dynamics to subsequent memory on a trial-by-trial basis, jackknife inter-trial phase coherence was computed from the lowest-p significant whole-head channel × time cluster for each band × effect combination identified in the group-level analyses (Richter et al., 2015). For each trial j, ITPC was recomputed after excluding that trial, within each participant and Selection × Feedback condition, pooling all memory outcomes:

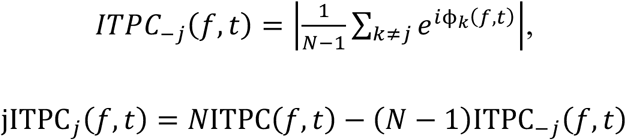

where ITPC(*f*, *t*) is the full-sample estimate from the corresponding participant and condition, and N is the number of trials in that condition. Pseudo-values were computed within conditions rather than within memory-outcome groups because ITPC is biased upwards in small samples and the jackknife does not remove this bias. Grouping by later memory would therefore give higher values to the smallest group, here Remember trials, regardless of any true association (see Text S1).

Each trial’s jITPC pseudo-value was averaged across the selected channels and frequency bins over the full temporal interval associated with the relevant cluster, then standardized within participant before model fitting. Once a cluster had been defined by the condition-level analysis, its jITPC values were extracted across all retained trials. The predictor was therefore computed without using the memory outcome it was then used to predict. Selection-related theta and alpha clusters each yielded middle-frontal and middle-posterior predictors. Feedback-related alpha yielded both regional predictors, whereas feedback-related theta yielded only a middle-posterior predictor. One additional predictor summarized the entire alpha Selection × Feedback interaction cluster. Regional subdivision was performed after whole-head inference and did not define the inferential search space. These eight jITPC predictors entered the primary joint model.

#### Response timing and latency sensitivity analyses

Response-initiation latency was defined as the interval between the stimulus-onset trigger and the response-start trigger preceding movement. Movement duration was taken from the behavioral task records. Condition summaries were based on participant-level medians over the 6,679 retained action-onset-locked epochs. For the condition-level ITPC sensitivity analysis, self-generated and externally generated trials were matched within each participant × feedback condition on log-transformed latency using a pooled-stratum caliper of .20 SD. Participants were retained when at least 15 matched pairs were available in both feedback (i.e., fluent vs. disrupted) conditions. This criterion retained 35 participants. ITPC was recomputed from the matched trials using the primary time-frequency settings, and self-generated–externally generated contrasts were tested using two-sided channel × time cluster-based permutation tests with 10,000 randomizations. Trial-wise neural-memory sensitivity models additionally included within-participant-standardized log latency and the standardized participant mean log latency.

#### Theta–alpha cross-frequency phase synchronization (CFS)

At each sensor, 2:1 theta–alpha phase synchronization was quantified from the phase difference between twice the theta phase and the alpha phase during the action-onset-locked 0–1-s interval, using 300-ms windows advanced in 50-ms steps. To control for phase locking attributable to the individual bands, 1,000 surrogate estimates were generated by disrupting theta–alpha trial correspondence within each participant and condition while preserving the theta and alpha time series. Surrogate-corrected coupling was defined as observed coupling minus the mean surrogate value. Raw coupling and surrogate-standardized z scores were examined as sensitivity measures. Selection, Feedback, and Selection × Feedback effects were evaluated using two-sided channel × time cluster-based permutation tests with a cluster-forming threshold of *p* < .05, at least one neighboring channel, and 10,000 Monte Carlo randomizations (Herweg et al., 2016; Siebenhühner et al., 2016).

#### Source-level theta and alpha phase alignment

Source activity was reconstructed in FieldTrip using the standard boundary-element head model and a 10-mm Montreal Neurological Institute (MNI) template source grid because individual MRI and digitized electrode coordinates were unavailable. The grid was restricted to cortical masks defined by the automated anatomical labeling (AAL) atlas. Source positions without an exact atlas match were assigned to the nearest labeled cortical voxel within 6 mm. Condition-independent common linearly constrained minimum variance (LCMV) filters were computed across all valid encoding trials using 5% regularization, fixed orientations, and unit-noise-gain weight normalization, and were applied identically to the four Selection × Feedback conditions. Trial-wise sensor data were band-pass filtered at 4–8 Hz for theta or 8–13 Hz for alpha, transformed to complex analytic signals using the Hilbert transform, and projected through the common filters. Source ITPC was calculated at each cortical source position from 0 to 1,000 ms after action onset. ITPC maps were averaged into 20-ms temporal bins only after trial-level phase estimation, and no trial-level temporal resampling was applied. Separate two-sided source-position × time cluster-based permutation tests evaluated the main effects of Selection and Feedback and their interaction. Clusters were formed at a sample-level threshold of *p* < .001 and tested at cluster-level *p* < .05 using 10,000 Monte Carlo randomizations (Xie et al., 2025; Li et al., 2026).

#### Anatomical localization of source-level results

The AAL atlas distributed with FieldTrip was used to anatomically label all reported source-level ITPC and PAC results, including theta-and alpha-ITPC clusters and data-driven PAC seed and target coordinates. For ITPC clusters, labels were obtained for the peak voxel and the absolute–*t*-weighted cluster center, and anatomical composition was audited across the unique spatial voxels comprising each cluster. For PAC analyses, the selected seed centers and reported target locations were labeled using the same atlas procedure. Some fixed literature-derived PAC seeds retained their prespecified coordinates and anatomical designations. Exact atlas matches were used where available, and otherwise, the nearest labeled atlas voxel within 6 mm was used. Atlas labels were used for anatomical description and did not determine cluster formation or statistical significance.

#### Source-level theta–gamma phase–amplitude coupling

To test whether agency-related encoding effects extended beyond local phase alignment to coordination across regions and frequencies, source-level theta–gamma PAC was estimated during the early encoding period (Tort et al., 2009; Heusser et al., 2016). Source reconstruction was performed in FieldTrip using a common broadband linearly constrained minimum variance (LCMV) beamformer filter computed across all valid encoding trials and then applied identically across conditions and frequency components (Li et al., 2026). The broadband filters inherited the same 10-mm source geometry and AAL-defined cortical mask used for the source-level ITPC analyses.

Three 10-mm theta-phase seeds were selected using leave-one-participant-out (LOSO) searches constrained to the significant source-level theta Selection cluster: right cuneus (MNI: 15, −90, 20), right posterior cingulate cortex (PCC; MNI: 5, −40, 20), and left calcarine cortex, encompassing primary visual cortex (V1; MNI: −5, −90, 0). Five literature-motivated sensitivity seeds were additionally examined: left and right inferior frontal gyrus (BA45) (MNI: −54, 21, 12 and 54, 21, 12), left and right pre-supplementary motor area (pre-SMA; MNI: −12, 26, 56 and 12, 26, 56), and dorsal anterior cingulate cortex (dACC; MNI: −5, 40, 20).

For the data-driven seed search, candidate theta-phase seed locations were restricted to voxels within the significant positive selection-related source-level theta-ITPC cluster. In each leave-one-participant-out (LOSO) fold, PAC effect maps from the held-out participant were omitted, and the remaining participants were used to select the seed, gamma-target component, and time window with the largest connected suprathreshold mass (sum of absolute *t* values; two-sided uncorrected *p* < .05). The mean PAC effect within this training-defined component was then extracted from the held-out participant. LOSO was repeated for every participant and separately for Selection, Feedback, and Selection × Feedback effects and for positive and negative effects. Because the upstream ITPC cluster was defined in the full sample, LOSO reduced circularity in PAC seed-target-window selection but did not make the initial candidate-seed mask fully independent of the data.

Theta phase (4–8 Hz) at each seed was coupled with gamma amplitude (30–80 Hz) at whole-cortex targets within the AAL-defined source mask during 0–1,000 ms action-onset-locked intervals, using 500-ms windows advanced in 100-ms steps. For each trial, seed-to-voxel PAC was quantified using amplitude-weighted mean vector length (MVL):

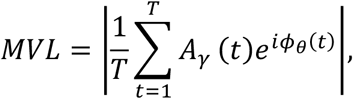

where *A_γ_*(*t*) is gamma-band amplitude at the target voxel, *φ_θ_*(*t*) is theta-band phase at the seed region, and *T* is the number of samples in the analysis window.

Normalized MVL was calculated by dividing raw MVL by the mean gamma amplitude at the corresponding target voxel within the same analysis window:

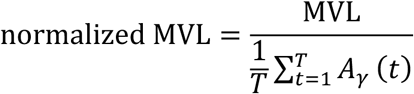

Normalized MVL was the primary coupling metric, whereas raw MVL was examined as a complementary sensitivity measure. The primary condition-level analyses tested Selection, Feedback, and Selection × Feedback separately using source-position × PAC-window time cluster-based permutation tests. For each effect family, *p* values were corrected across the 8 theta seeds using BH–FDR correction.

For the two FDR-significant normalized MVL interaction clusters, a targeted within-condition trial-shuffled surrogate analysis was conducted as a post-selection sensitivity analysis. Gamma-amplitude trial series were paired with theta-phase trial series from different trials while preserving the within-condition phase and amplitude distributions. The resulting null quantified PAC expected after disrupting trial-wise phase-amplitude correspondence. It was not treated as an independent replication or as the normalization procedure itself.

#### Exploratory source-level alpha–gamma coupling

Alpha-phase seeds were selected using the same leave-one-participant-out seed-search principle as in the theta analysis. Coupling between alpha phase (8–13 Hz) and gamma amplitude (30–80 Hz) was quantified using normalized MVL. Selection, Feedback, and Selection × Feedback were treated as separate effect families, with BH–FDR correction across the five alpha seeds within each family. For the five FDR-retained feedback-related clusters, a targeted sensitivity analysis used 1,000 within-condition trial derangements to disrupt alpha-phase/gamma-amplitude trial correspondence. Group inference tested participant-level observed-minus-mean-surrogate effects with two-sided sign-flip tests and BH–FDR correction across the five frozen targets. Because targets were selected from the primary exploratory results, these tests were interpreted as post-selection sensitivity analyses rather than independent confirmation.

#### Trial-wise phase–amplitude coupling predictors

Trial-wise jPAC predictors were derived from the two significant normalized-MVL Selection × Feedback interaction clusters using leave-one-trial-out jackknife estimates (Richter et al., 2015). Each predictor used the theta seed, gamma-target voxels, and time window defined by its condition-level cluster. For each trial j, a leave-one-out PAC estimate was computed after excluding that trial, within each participant and Selection × Feedback condition, pooling all memory outcomes:

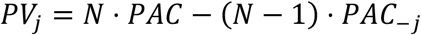

where *N* is the number of trials in that condition, *PAC* is the full-sample coupling estimate, and *PAC*_−j_ is the leave-one-out estimate obtained after removing trial *j*. Each predictor was averaged within the cluster-defined target voxels and time window, extracted across all retained trials, and standardized within participant. The two normalized jPAC predictors were entered jointly with the 8 channel-level jITPC predictors in the primary memory model. Raw-MVL analyses were treated as sensitivity analyses.

To test whether PAC–memory associations were reducible to target-region gamma amplitude, follow-up models included gamma amplitude extracted from the same source-level target cluster.

### Trial-wise prediction analyses (Experiment 3)

#### Bayesian mixed-effects modeling of trial-wise neural predictors

Bayesian generalized linear mixed-effects models were fitted in R (version 4.5.1) using brms (version 2.22.0). Neural predictors were derived from significant condition-level clusters, extracted across all retained trials, and standardized within participant. The trial-wise neural–memory analyses included 4,994 trials from 42 participants. A multinomial model predicted Remember, Know, and Forgotten responses, with Forgotten as the reference category. The primary joint model included all eight jITPC predictors and both normalized jPAC predictors, together with Selection, Feedback, and their interaction. Remember-versus-Know contrasts were calculated for each posterior draw by subtracting the Know-versus-Forgotten coefficient from the Remember-versus-Forgotten coefficient. Bernoulli-logit models predicted correct versus incorrect recognition of the unselected item using the same 10 neural predictors. Follow-up PAC-control models additionally included gamma amplitude from the corresponding source-level target cluster to assess whether PAC–memory associations were independent of regional gamma– amplitude variation.

Sense of agency ratings and time-compression estimates were not included as covariates in the memory models. They were analyzed separately in Gaussian mixed-effects models with Selection, Feedback and their interaction as fixed effects.

Across multinomial, unselected-item, and Gaussian models, Selection and Feedback were sum coded. Participant was modeled with varying intercepts and varying slopes for Selection, Feedback, and their interaction. An additional varying intercept was included for each encoding trial (Item) in all three model types.

Sampling settings and priors for these trial-wise models followed those described under Behavioral measures and statistical analyses. For the multinomial memory model, multiplicity was controlled across the 30 prespecified neural predictor-by-memory contrasts using Bayesian aggregate false-sign-rate correction (Stephens, 2017). Local false-sign rates were ordered and cumulatively averaged to obtain *q*_BFSR_ values. Neural effects were considered robust when their 95% credible intervals excluded zero and *q*_BFSR_ < .05. Multiplicity for unselected-item recognition was controlled separately across the same ten primary neural predictors.

#### Cross-participant decoding

Cross-participant predictive utility of trial-wise neural features was evaluated in Python 3.13.5 with scikit-learn 1.6.1. All analyses used leave-one-participant-out cross-validation across 42 participants and a fixed L1-penalized logistic-regression classifier (liblinear, C = 1, class-weight balanced) without inner-loop hyperparameter search. Predictors were standardized using parameters estimated within each training fold and then applied to the held-out participant. This follow-up focused on early middle-frontal alpha jITPC (127–245 ms) and later middle-posterior theta jITPC (353–570 ms), selected following the neural–memory association analyses. Neural predictors were computed within participant × Selection × Feedback groups, without using subsequent memory labels. Six feature sets were evaluated: experimental conditions alone (Selection and Feedback), alpha alone, theta alone, alpha plus conditions, theta plus conditions, and both neural predictors plus conditions. Five classification tasks were tested: three-class Remember/Know/Forgotten classification, Remember versus Forgotten, Know versus Forgotten, Remember versus Know, and correct versus incorrect unselected-item recognition. Performance was quantified using pooled out-of-fold area under the receiver operating characteristic curve (AUC). Three-class classification used a one-versus-rest approach, summarized by macro-average AUC and category-specific one-versus-rest AUCs. Confidence intervals were estimated from 10,000 participant-level bootstrap samples of the saved out-of-fold predictions, without refitting the models. Paired comparisons used identical participant resamples to estimate AUC differences between models and between memory categories.

#### Focused moderation analysis

This exploratory analysis addressed the preregistered agency-moderation hypothesis, by testing whether explicit agency moderated neural associations with Remember versus Know responses. Binary logistic generalized linear models were fitted with participant-clustered robust standard errors. Each model included one neural predictor, explicit agency, their interaction, and Selection and Feedback as covariates. Neural predictors were computed within participant × Selection × Feedback groups without using subsequent memory labels. Neural predictors and explicit agency ratings were standardized within participant. Interaction-term p values were adjusted using the BH–FDR procedure across the 10 neural predictor × explicit agency tests for the Remember-versus-Know contrast, comprising eight jITPC and two jPAC predictors.

### Data availability

The data and analysis scripts underlying the findings are available on the Open Science Framework at https://osf.io/3yhpz. The preregistration is available at https://doi.org/10.17605/OSF.IO/Z43DF.

### Code accessibility

Custom scripts for behavioral and EEG analyses, statistical modeling, and figure generation are available through the Open Science Framework project at https://osf.io/3yhpz.

## Results

The three experiments shared a 2 × 2 encoding design crossing action selection (self-generated vs externally generated) with action feedback (fluent vs disrupted) (Fig. 1). The behavioral results from all three experiments are reported first and together, followed by the EEG results from Experiment 3.

**Figure 1.**
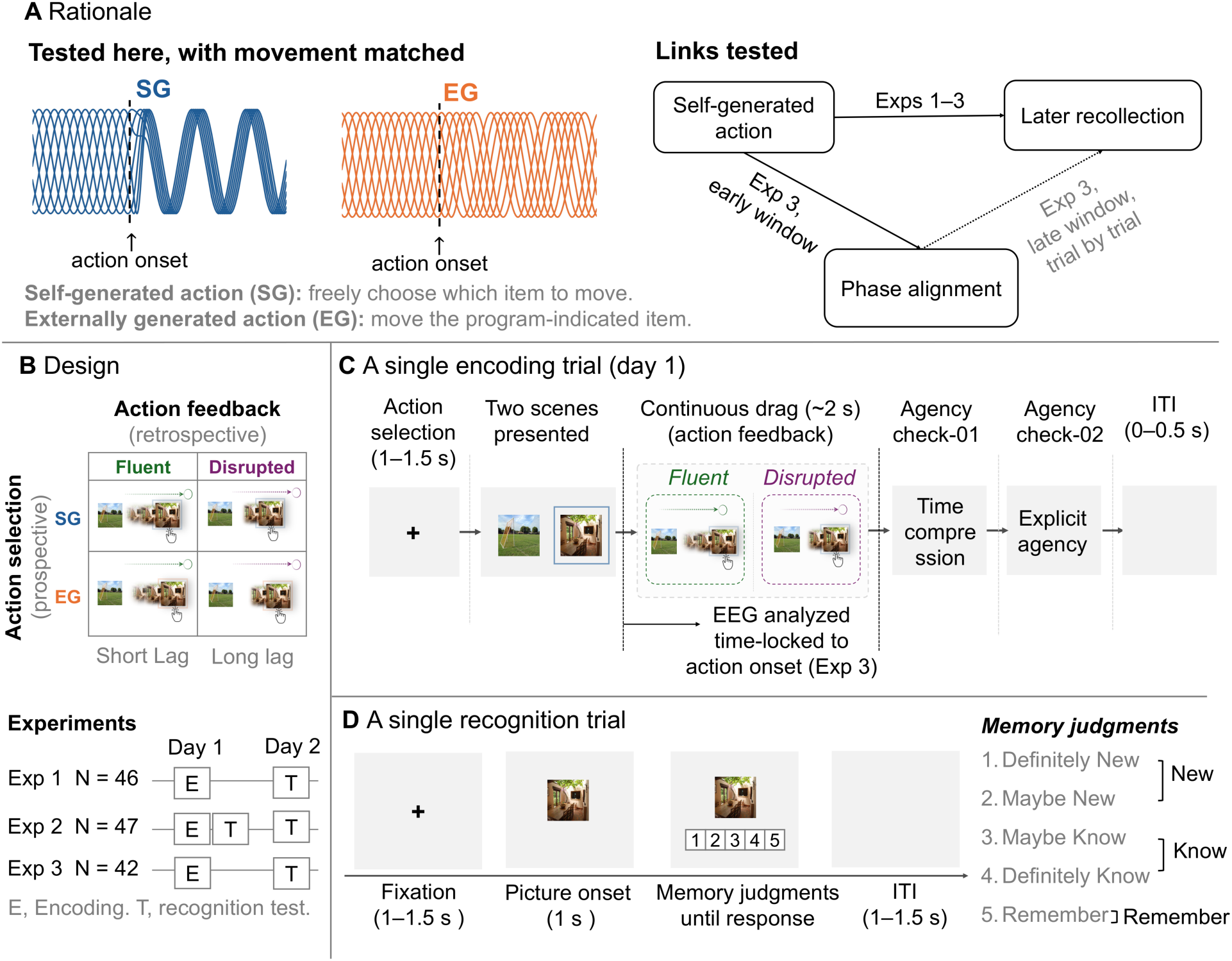
Rationale and experimental design. **A**, Rationale. The schematic traces illustrate the proposed increase in phase alignment after self-generated action, with movement matched. These traces are illustrative and do not represent observed data. SG denotes freely choosing which item to move, and EG moving the item indicated by the program. The diagram summarizes the proposed links between action selection, phase alignment at encoding, and later recollection. Solid arrows mark effects of the manipulation, and the dotted line marks a trial-wise association. Experiment 3 examined condition-level phase alignment and its trial-wise association with later memory. **B**, Design. The within-participant 2 × 2 design crossed action selection (self-generated, SG, or externally generated, EG) with action feedback (fluent or disrupted). Feedback was set by the image-following coefficient during dragging (.09 fluent, .01 disrupted), so that at each update the image moved 9% or 1% of the remaining distance to the mouse position. Each encoding trial yielded a time-compression estimate and an explicit agency rating. The final analyzed samples comprised 46, 47, and 42 participants in Experiments 1–3, respectively. E and T denote encoding and recognition test. **C**, A single encoding trial. In the encoding phase, a 1–1.5 s selection cue was followed by two scene images. Participants dragged the self-selected or program-selected image for approximately 2 s under the assigned feedback, then gave a time-compression estimate and an explicit agency rating. A 0–0.5-s inter-trial interval followed. EEG was recorded during encoding in Experiment 3 and analyzed relative to action onset. **D**, A single recognition trial. In the recognition phase, a single scene image appeared at the center of the screen, and participants judged it new, familiar without specific detail (Know), or remembered with specific detail from the encoding episode (Remember). The five response options (Definitely New, Maybe New, Maybe Know, Definitely Know, and Remember) were grouped into New, Know, and Remember categories as shown. Test items were selected and unselected scenes from encoding and new scenes. Recognition was tested about 24 h after encoding (Experiments 1 and 3) or both immediately and after about 24 h (Experiment 2). EEG, electroencephalography; ITI, inter-trial interval.

### Self-generated action benefited recollection most and, more weakly, the unselected item

The three experiments showed a consistent pattern. Self-generated action increased Know and Remember responses, Remember most, and to a lesser extent improved memory for the unselected scene (Fig. 2).

**Figure 2.**
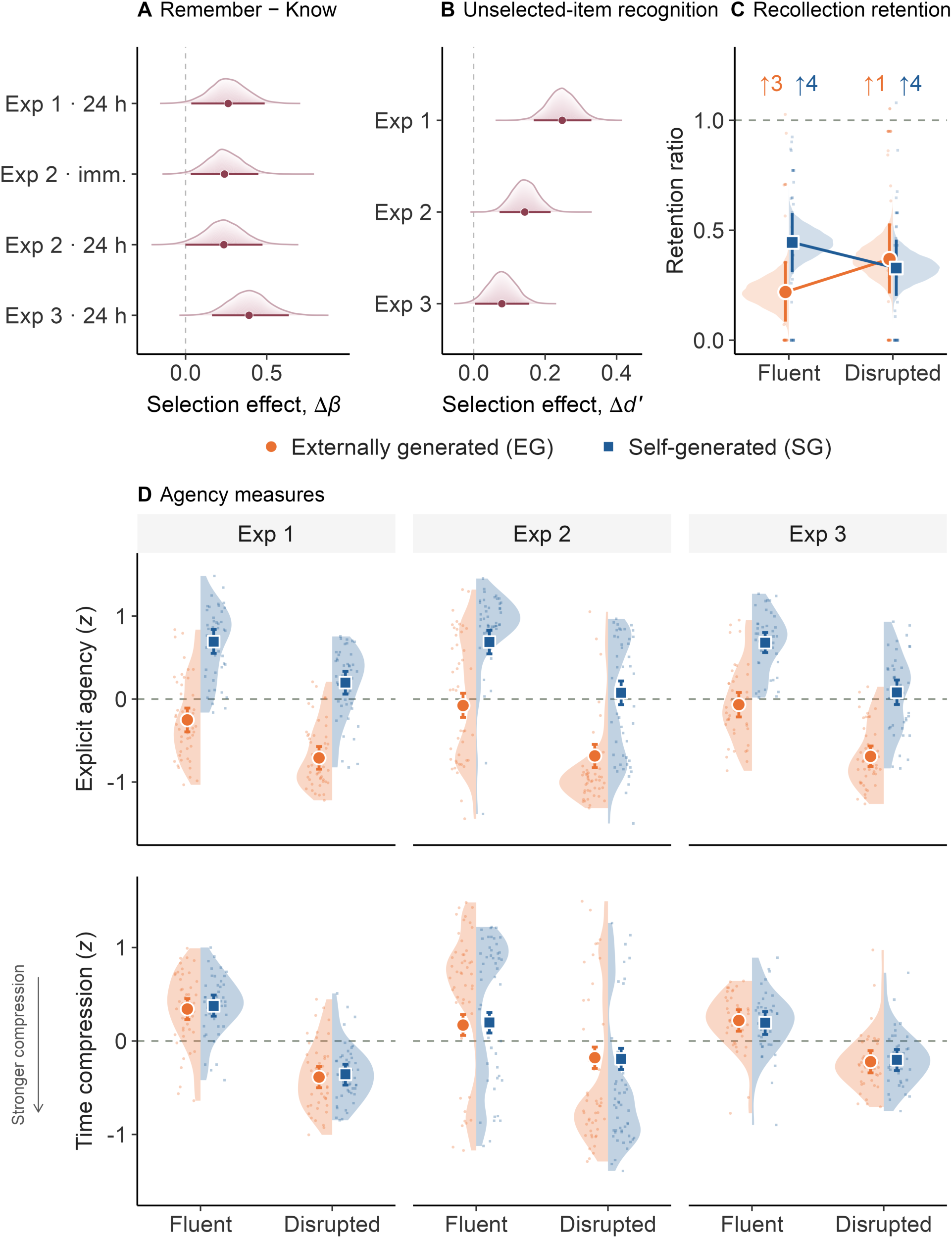
Self-generated action benefited recollection more than familiarity and, more weakly, the unselected item. **A**, Posterior distributions of the difference between the Selection coefficients for Remember and Know responses (Δ*β* = *β_Remember_* – *β_Kno_*_w_) for selected items in Experiments 1–3. Experiment 2 includes both immediate and 24-hour tests. Experiments 1 and 3 include the 24-hour test only. Positive values indicate a larger Selection effect on Remember than Know responses. **B**, Posterior differences in discriminability (*d′*) for the unselected, co-present item after self-generated versus externally generated action in Experiments 1–3. Positive values indicate a memory advantage after self-generated action. **C**, Recollection in Experiment 2 across the immediate and 24-hour tests, expressed as the delayed-to-immediate ratio. The dashed horizontal line indicates equal performance at the two tests. Arrow annotations give the number of participants with ratios above 1.08. These values were not plotted but were included in the analyses. **D**, Standardized explicit agency ratings and time-compression estimates across the four Selection × Feedback conditions in Experiments 1–3. More negative time-compression values indicate stronger perceived temporal compression. Gray dashed horizontal lines mark zero on the standardized scales. In **A** and **B**, shaded densities show posterior distributions, points indicate posterior means, and horizontal intervals indicate 95% credible intervals. In **C**, shaded densities show posterior distributions, and in **D**, split violins show participant-level distributions. In **C** and **D**, large symbols indicate posterior means, vertical intervals indicate 95% credible intervals (CrIs), and small semi-transparent points show participant-level estimates. In **C** and **D**, orange denotes externally generated action, and blue denotes self-generated action. Fluent and disrupted feedback had shorter and longer lags between hand and scene movement. SG, self-generated; EG, externally generated.

#### Recollection more than familiarity

In Experiment 1, a Bayesian multinomial model showed that self-generated relative to externally generated action increased Know responses (*β* = .23, 95% CrI [.11, .34]) and Remember responses (*β* = .49, 95% CrI [.28, .69]), with the posterior mean coefficient for Remember responses approximately twice that for Know responses. Disrupted feedback had no credible effect on familiarity (*β* = .11, 95% CrI [-.01, .22]) or on recollection, and the Selection × Feedback interaction was not credible for either outcome. The selection benefit was therefore present for familiarity and larger for recollection (Fig. 2A). Experiment 3 replicated this pattern (Know *β* = .22, 95% CrI [.12, .33]; Remember *β* = .61, 95% CrI [.40, .84]). The Remember effect held after adjustment for the longer response-initiation latency of self-generated trials (*β* = .55, 95% CrI [.35, .77], Text S1). In Experiment 2, the direct Remember-versus-Know contrast also favored self-generated action at the immediate test (*β* = .24, 95% CrI [.03, .45], Text S1). Confidence-ROC model comparisons provided criterion-dependent support for a dual-process account of recognition (Text S1) (Yonelinas, 1994; Wixted, 2007; Yonelinas and Parks, 2007). In hierarchical drift-diffusion models of recognition, self-generated action raised the drift rate for both Remember and Know responses in Experiments 1 and 3, consistent with stronger evidence accumulation for selected-item recognition (Fig. S3 in Text S1).

#### Recognition of the unselected item

If the benefit reflected attention paid to the chosen item alone, only that item should improve. Memory for the co-present, unselected item tested this. In Experiment 1, recognition was higher after self-generated than externally generated action (*β* = .25, 95% CrI [.12, .39]), with no credible Selection × Feedback interaction (Fig. 2B). In Experiment 2, the benefit was clearest at the immediate test and uncertain after 24 hours. Experiment 3 showed a small advantage in discriminability (unselected-item Δ*d′* = .08, 95% CrI [.003, .16]), but the trial-wise Selection effect was uncertain both before and after adjustment for response-initiation latency (adjusted *β* = .04, 95% CrI [-.07, .15]). After adjustment, the benefit remained in Experiment 1 (*β* = .23, 95% CrI [.10, .38]) and at the immediate test in Experiment 2 (*β* = .19, 95% CrI [.05, .33]; Text S1). The benefit was therefore not confined to the acted-upon image, so a preference for the chosen scene alone cannot fully explain it. Choosing between two scenes requires evaluating both, which could raise attention to the scene not chosen. Attention was not measured, so the design does not separate that account from an event-level effect of agency (see Discussion).

#### Immediate versus 24-hour test

Experiments 1 and 3 tested memory after 24 hours. Experiment 2 added an immediate test to the same design, so the benefit could be compared across the two tests. Recollection declined more than familiarity over the 24 hours (Text S1). How much it declined depended on selection and feedback together: recollection retention, the ratio of delayed to immediate performance, showed a credible Selection × Feedback interaction (*β* = −.07, 95% CrI [-.13, −.01]).

Under fluent feedback, recollection declined less after self-generated than externally generated action (Δ*M* = .23, 95% CrI [.07, .38]). Under disrupted feedback, it did not (Δ*M* = −.04, 95% CrI [-.21, .12]), a credible difference-in-differences (disrupted minus fluent: −.27, 95% CrI [-.50, −.04]). This Selection × Feedback interaction was larger for recollection than for familiarity (Δ*M* = .27, 95% CrI [.04, .50]) (Fig. 2C).

#### Explicit agency and time compression

Explicit agency ratings were higher for self-generated than externally generated action in every experiment (Exp 1 *β* = .46, 95% CrI [.35, .57]; Exp 2 *β* = .38, 95% CrI [.27, .50]; Exp 3 *β* = .38, 95% CrI [.28, .47]) and higher under fluent than disrupted feedback. The time-compression index, the difference between estimated and actual image-movement duration, varied with action feedback, with more negative values (stronger compression) under disrupted than fluent feedback, but not with selection (Fig. 2D). Explicit agency thus tracked both components, and time compression only the outcome-related one, consistent with separable prospective and retrospective components of agency. Time compression is an indirect index, so it is interpreted cautiously.

### Selection and feedback altered phase alignment at different latencies

Behavioral findings show what was remembered but not how encoding carried the advantage. Experiment 3 therefore recorded EEG during encoding. Analyses were time-locked to action onset, when selection had been made and feedback began. Movement duration was about 2.06 s in every condition (Text S1), so differences in movement duration are unlikely to explain the effects.

Phase alignment was measured as action-onset-locked inter-trial phase coherence (ITPC), the consistency of oscillatory phase across trials (Long et al., 2025). ITPC was higher for self-generated than externally generated action in theta across a widespread fronto-posterior cluster from 19–416 ms (*p*_cluster_ < .001; *q*_FDR_ < .001), and in alpha, in a more focal cluster from 127–245 ms (*p*_cluster_ < .001; *q*_FDR_ < .001). Disrupted feedback lowered ITPC later, in posterior theta (353– 570 ms; *p*_cluster_ < .001; *q*_FDR_ < .001) and in frontal and posterior alpha (301–590 ms; *p*_cluster_ < .001; *q*_FDR_ < .001) (Fig. 3A–D). Self-generated action therefore raised earlier alignment, and disrupted feedback lowered later alignment. A Selection × Feedback interaction appeared in a posterior alpha-band cluster from 145–282 ms (*p*_cluster_ < .001; *q*_FDR_ < .001). Shorter interaction clusters (< 50 ms) were not interpreted. Self-generated trials had longer response-initiation latencies than externally generated trials (participant-level medians about 1.0 and 0.7 s). Both selection effects remained significant after self-generated and externally generated trials were matched on latency (latency-matched theta cluster 48–373 ms, peak POz *t* = 8.27, *p*_cluster_ < .001, *q*_FDR_ < .001; alpha 134– 209 ms, peak PO3 *t* = 5.13, *p*_cluster_ < .001, *q*_FDR_ < .001; Text S1), so the longer latency is unlikely to explain them.

**Figure 3.**
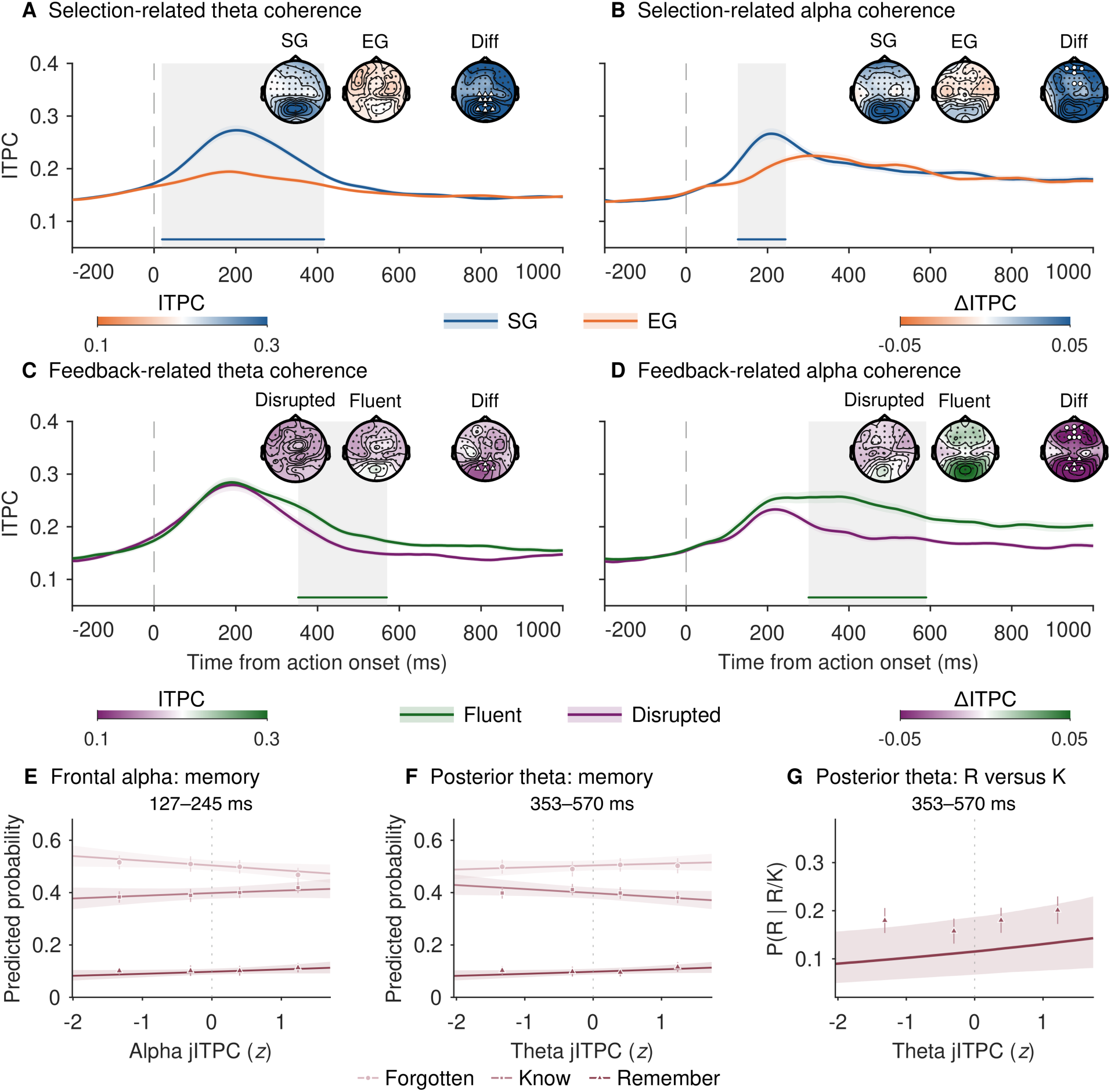
Selection and feedback altered theta and alpha phase alignment at different latencies. Action-onset-locked theta and alpha inter-trial phase coherence (ITPC) during encoding in Experiment 3. **A** and **B**, Effects of self-generated versus externally generated action in the theta (**A**; 4–7.99 Hz) and alpha (**B**; 8–12.99 Hz) bands. Frequency ranges denote nominal band boundaries, and the corresponding sampled frequency bins are specified in Methods. **C** and **D**, Effects of disrupted versus fluent feedback in the theta (**C**) and alpha (**D**) bands. Selection-related time courses are averaged across feedback conditions, and feedback-related time courses are averaged across selection conditions. In the time courses, blue and orange denote self-generated and externally generated action in A/B, and green and purple denote fluent and disrupted feedback in C/D. Lines indicate means across participants and ribbons indicate ±1 SEM (n = 42). Gray shading and colored horizontal bars mark the significant cluster intervals: 19–416 ms (**A**), 127–245 ms (**B**), 353–570 ms (**C**), and 301–590 ms (**D**). The vertical line denotes action onset. Scalp maps show condition-averaged ITPC and the corresponding contrast averaged across each cluster interval. Difference maps represent self-generated minus externally generated action in **A** and **B** and disrupted minus fluent feedback in **C** and **D**. Condition maps use orange-to-blue scales in A/B and purple-to-green scales in C/D. Difference maps use orange-white-blue scales in A/B and purple-white-green scales in C/D, with white indicating zero. Outlined markers identify the middle-frontal and middle-posterior channel groups subsequently used to construct trial-wise predictors. **E**–**G**, Trial-wise associations between jackknife ITPC (jITPC) and subsequent memory. Middle-frontal alpha jITPC from the selection cluster (127– 245 ms) was positively associated with Remember versus Forgotten responses, whereas its association with Know versus Forgotten responses was uncertain (**E**). Middle-posterior theta jITPC from the feedback cluster (353–570 ms) showed an uncertain association with Remember versus Forgotten responses (**F**) and a positive association with Remember versus Know responses (**G**). Lines show posterior predictions (random effects at zero), and ribbons show 95% credible intervals. Points show participant-balanced descriptive proportions. In G, they lie above the line because, at low probabilities, the average of participant probabilities exceeds the probability at the average log-odds. Triangles, squares, and circles denote Remember, Know, and Forgotten responses, respectively. SG, self-generated; EG, externally generated. Fluent and disrupted denote fluent and disrupted action feedback, respectively.

### Trial by trial, later posterior theta alignment distinguished recollection from familiarity

The condition-level effects showed that selection and feedback changed phase alignment. The trial-wise analysis asked whether alignment also predicted what was later remembered. Jackknife ITPC (jITPC) gave a trial-wise estimate of alignment for each significant cluster (Richter et al., 2015). Estimates were computed within each participant and Selection × Feedback condition without reference to later memory, avoiding bias arising from unequal trial counts across subsequent-memory groups (Text S1). These estimates entered a joint Bayesian multinomial model with false-sign-rate control (Fig. 3E–G). Middle-posterior theta alignment in the later feedback window (353–570 ms) was higher on trials later given Remember than Know responses (*β* = .14, 95% CrI [.01, .27]; *q*_BFSR_ = .017) but did not distinguish Remember from Forgotten (*β* = .08, 95% CrI [-.04, .21]). Middle-frontal alpha alignment in the selection window (127–245 ms) predicted Remember over Forgotten (*β* = .15, 95% CrI [.006, .29]; *q*_BFSR_ = .019) but not Know over Forgotten or Remember over Know. No other alignment predictor showed a robust association (Text S1).

ITPC measures phase consistency across trials at one site and frequency. PAC measures how the amplitude of faster activity varies with the phase of slower activity. Because theta-organized gamma activity (30–80 Hz) has been linked to episodic memory (Tort et al., 2009; Nyhus and Curran, 2010; Heusser et al., 2016), trial-wise theta–gamma PAC in source-reconstructed signals provided a further test of the binding account. Neither coupling predictor, calcarine–frontal or posterior-cingulate–cortical, showed a clear association with any memory contrast (Text S1).

Source-space PAC from scalp EEG is limited by source-localization uncertainty and possible ocular or muscle contamination of gamma amplitude, so this null result is weak evidence against a role for coupling. The full source analysis includes source-level theta and alpha ITPC (Fig. S1), condition-level PAC (Fig. S2), the eight-seed search, the surrogate, raw-metric and gamma-amplitude controls, and an exploratory alpha–gamma PAC analysis (Text S1).

### Phase alignment added modest predictive information across participants

Exploratory leave-one-participant-out decoding tested whether phase alignment added predictive information beyond Selection and Feedback. The two neural predictors were chosen after the association analyses (see Materials and Methods). Adding frontal alpha increased Remember-versus-Forgotten AUC from .588 to .603 (ΔAUC = .014, 95% CI [.007, .022]), and adding posterior theta increased Remember-versus-Know AUC from .532 to .556 (ΔAUC = .024, 95% CI [.009, .039]). Posterior theta also improved Remember-versus-Forgotten decoding (ΔAUC = .019, 95% CI [.009, .029]). Both neural predictors also improved Know-versus-Forgotten decoding and, by smaller amounts, unselected-item decoding, whereas alpha did not improve Remember-versus-Know decoding (Fig. 4A). The alpha-plus-condition Remember-versus-Forgotten model and the theta-plus-condition Remember-versus-Know model both had bootstrap confidence intervals above the chance AUC of .50 (alpha, 95% CI [.571, .637]; theta, 95% CI [.525, .587], respectively), although discrimination remained modest. In the three-class model combining alpha, theta and the experimental conditions, one-versus-rest AUC was higher for Remember than Know responses (.576 versus .524; paired ΔAUC = .052, 95% CI [.015, .089]), but this advantage was also present in the condition-only model (Fig. 4B). Phase alignment therefore added a small amount of predictive information that generalized to held-out participants, and only theta added information that separated recollection from familiarity.

**Figure 4.**
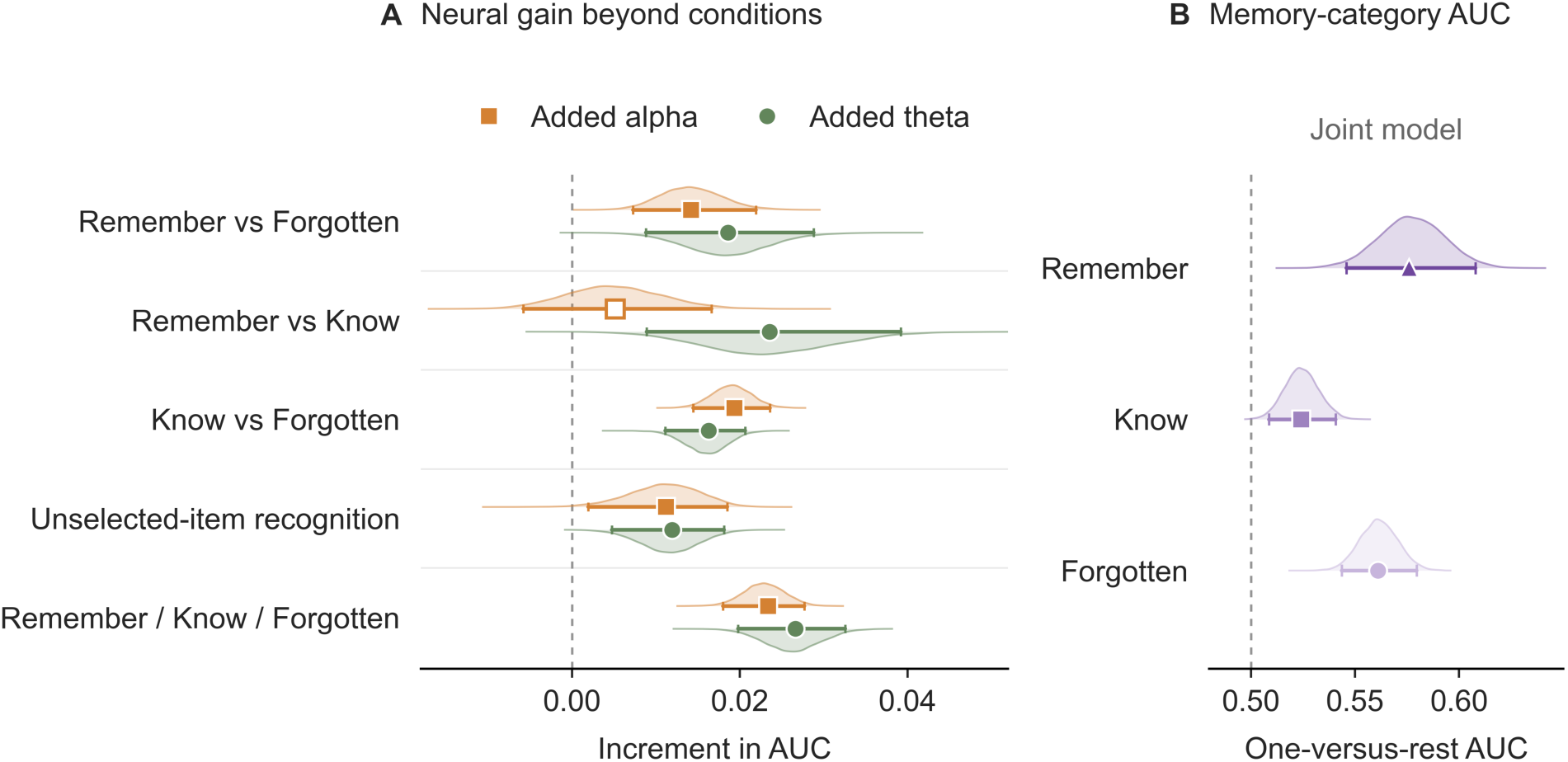
Decoding of later memory from trial-wise phase alignment. **A**, Increment in pooled out-of-fold AUC relative to a condition-only model containing Selection and Feedback. Orange squares indicate the addition of early middle-frontal alpha jITPC (127– 245 ms), and green circles indicate the addition of later middle-posterior theta jITPC (353–570 ms). Three-class performance is summarized by macro-averaged one-versus-rest AUC. **B**, Category-specific one-versus-rest AUCs from the three-class model containing alpha, theta, Selection, and Feedback. Remember, Know, and Forgotten responses are represented by dark-purple triangles, medium-purple squares, and light-purple circles, respectively. Each category is compared with the other two categories combined. In **A** and **B**, points show observed estimates, and horizontal intervals show percentile 95% confidence intervals from 10,000 participant-level bootstrap samples of the saved out-of-fold predictions. Pale half-violins show smoothed bootstrap distributions, scaled to equal peak height. In **A**, hollow markers indicate intervals containing zero, and filled markers indicate intervals excluding zero. Dashed lines indicate zero increment in **A** and chance AUC (.50) in **B**. Models used leave-one-participant-out cross-validation across 42 participants. These follow-up analyses were exploratory, and confidence intervals were not adjusted for multiple comparisons.

An exploratory analysis tested whether explicit agency moderated these associations. The interaction between selection-related theta alignment and explicit agency in predicting Remember-versus-Know responses was positive (*β* = .14, 95% CI [.05, .23]; *p* = .003). It survived BH–FDR correction across the 10 neural predictors × explicit agency interaction tests for this contrast (*q*_FDR_ = .028) (Fig. 5).

**Figure 5.**
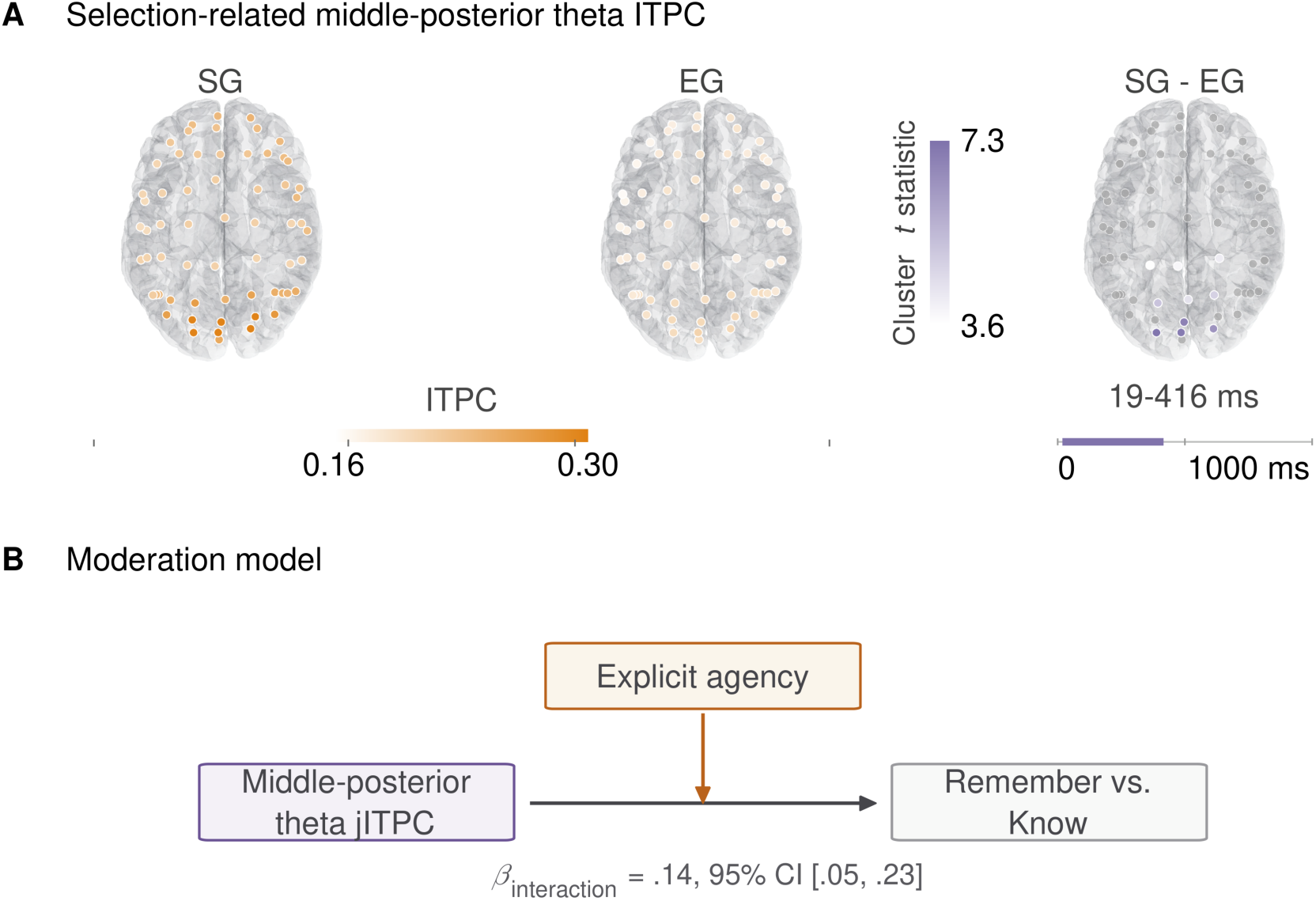
Exploratory analysis of explicit agency and the theta–memory association. Related to Fig. 3. **A**, Spatial characterization of the selection-related middle-posterior theta predictor. The first two maps show mean theta ITPC for SG and EG trials, and the SG–EG map shows cluster statistics for the channels used to construct the predictor. Colored markers indicate the predictor channels, whereas gray markers provide spatial context. The cortical surfaces are anatomical references and do not represent source-localized activity. **B**, Schematic of the moderation model. The reported coefficient and 95% confidence interval describe the interaction between selection-related theta jITPC and explicit agency. jITPC, jackknife inter-trial phase coherence; SG, self-generated; EG, externally generated.

## Discussion

Recollection is the form of memory in which the past is re-experienced as one’s own. Being the agent of an action at encoding shaped this form of memory. In three experiments, choosing which scene to act on improved later memory for that scene, more for recollection than for familiarity. Agency varied only at encoding, movement was matched, and the memory test was the same for all items. The findings therefore link encoding-stage agency to subsequent memory, without isolating the contributions of consolidation and retrieval. The pattern replicated across experiments, and in Experiment 3 it held after adjustment for the longer time taken to choose.

The first question was which part of agency matters and which memory benefits. Performing the action was identical across conditions, so it cannot explain the difference in memory. Disrupted feedback lowered explicit agency but had no credible effect on memory at 24 hours in Experiment 1. Memory instead followed selection, as the choosing account predicts. Wegner proposed that the feeling of doing marks actions as one’s own in memory (Wegner, 2004). In the theory of apparent mental causation (Wegner and Wheatley, 1999), selection here varied exclusivity (whether the program offered another cause of the action), and feedback varied consistency (whether the outcome matched the movement). Both manipulations affected rated agency, but their effects on memory did not simply follow the pattern of agency ratings. The feeling of doing alone therefore did not determine what was remembered. Explicit agency tracked both manipulations, whereas time compression tracked only feedback, which here changed outcome timing, consistent with prospective and retrospective signals that are integrated in explicit judgments but separable (Synofzik et al., 2008; Sidarus et al., 2013; Dewey et al., 2014). Dual-process accounts hold that recollection depends on binding the elements of an episode, whereas familiarity reflects graded strength (Yonelinas, 1994, 2001, 2002; Yonelinas and Jacoby, 1995), so the larger effect on recollection suggests that choosing strengthens binding. Confidence-ROC comparisons gave criterion-dependent support for this account, so a strength-based account cannot be fully excluded (Text S1). Items from all encoding conditions were tested together against the same new items, so differences between conditions are unlikely to reflect a shift in response criterion at test.

The first question also asked whether the benefit extends beyond the chosen item. The co-present item, neither selected nor moved, was better recognized after self-generated action in Experiment 1 and at the immediate test in Experiment 2, with a small advantage in discriminability in Experiment 3. An event-level effect of agency predicts this benefit, consistent with Conway’s view that the working self shapes what enters the episodic record (Conway and Pleydell-Pearce, 2000; Conway et al., 2004; Conway, 2005) and with the view that the episode is the unit of autonoetic experience (Tulving, 1983, 1985; Wheeler et al., 1997). The unselected-item benefit remained after latency adjustment in Experiment 1 and at the immediate test in Experiment 2. In Experiment 3, the selected-item benefit remained after latency adjustment, whereas the adjusted unselected-item effect was uncertain. The extra time in self-generated trials may reflect the additional evaluation involved in choosing between the two scenes. The benefit to the unselected item may therefore arise from this evaluation, a part of selecting, without requiring an effect of agency on the event as a whole. Tracking gaze during the choice, or a yoked condition in which the program’s choice appears after each participant’s typical choice time, would separate the two.

The second question was whether the benefit changes over a day. In Experiment 2, the selection benefit for recollection was already present at the immediate test, consistent with an effect at encoding. What changed over the day depended on feedback. Under fluent feedback, recollection declined less over the day after self-generated than externally generated action, and under disrupted feedback this advantage was absent. Choosing created the benefit, and control over the outcome influenced how well it lasted. This points to processes after encoding, such as consolidation, as well as to encoding itself.

The third question was whether agency leaves a trace in neural dynamics at encoding. Self-generated action raised theta and alpha phase alignment early after action onset, and disrupted feedback lowered alignment later. The selection effects held when trials were matched on response-initiation latency. Trial-wise associations with memory were modest. Later posterior theta alignment, in the window that feedback affected, was higher before Remember than Know responses, whereas its association with Remember versus Forgotten responses remained uncertain in the joint model. In the exploratory decoding, both alpha and theta improved Remember-versus-Forgotten classification, whereas only theta showed a clear gain for Remember-versus-Know classification. These findings link later theta alignment to the distinction between recollection and familiarity, consistent with theta phase supporting the binding that distinguishes the two (Guderian and Düzel, 2005; Clouter et al., 2017). An attention account predicts that alignment tracks recognition broadly. Frontal alpha alignment predicted Remember over Forgotten but not Know over Forgotten in the joint model, although adding it improved the decoding of both, so the data do not clearly favor either account for alpha. Theta– gamma PAC, however, showed no clear trial-wise association with subsequent memory.

Later posterior theta alignment showed an overall Remember–Know association, whereas in an exploratory analysis, the Remember–Know association for earlier selection-related theta alignment became more positive as explicit agency increased. Alignment near action onset may provide a temporal reference for encoding, and later alignment may support the integration of information as the action and its feedback unfold (Jutras et al., 2013; Backus et al., 2016; Clouter et al., 2017). The moderation pattern raises the possibility that the memory relevance of earlier alignment varies with the accompanying experience of control. However, this analysis was exploratory, and the association does not establish that subjective agency causally controls the theta–memory relationship. Alignment rose within theta and alpha, whereas surrogate-corrected theta–alpha phase synchronization was lower (Text S1), so choosing made each rhythm more consistent without coupling the two more strongly.

Together, the results suggest a two-component account of how agency enters memory. Choosing what to act on, the prospective component, strengthens encoding, most for recollection. Control over the outcome, the retrospective component, does not create the benefit but helps recollection of what was chosen to last. The account makes two predictions. With other materials and responses, choice benefits should still be largest for recollection. Disrupting control over the outcome after a choice should reduce delayed more than immediate recollection.

These findings bear on two literatures. For memory research, the results show which part of acting benefits memory. Choosing improves recognition of the chosen item (Murty et al., 2015; DuBrow et al., 2019), and active learning outperforms passive learning (Markant et al., 2016), but in active learning, choosing, moving and controlling the outcome usually occur together. With movement held constant, choosing alone improved memory, most for recollection, and control over the outcome affected how well the benefit lasted. Active-learning advantages may therefore depend in part on selection, a possibility that can be tested in other tasks by holding movement constant. For research on the self, agency is a feature of the minimal, experiential self, and autonoetic recollection a feature of the self extended in subjective time (Gallagher, 2000; Tulving, 2002). The results suggest a link between the two: the self that chooses may shape what is later re-experienced. Remember judgments index reported recollection rather than the sense that a memory is one’s own, so this link is indirect. Self-referential encoding also strengthens recollection more than familiarity (Conway and Dewhurst, 1995; Lawrence and Chai, 2021), and self-relevance has been proposed to prioritize what enters cognition (Cunningham et al., 2011; Estudillo and Bindemann, 2014; Sui and Humphreys, 2015). Agency concerns authorship of an action, and self-reference the relation of information to self-knowledge, so the two are not equated here. Whether they share a route into memory remains to be tested.

Three limitations concern the neural results. First, neural–memory associations are correlational, so a causal role for theta alignment would need an intervention, such as rhythmic stimulation during encoding. Second, scalp phase measures are coarse. The early and late windows overlapped and arose from different contrasts, with wavelet smoothing limiting temporal precision. Early estimates may include pre-action activity, and phase resetting cannot be separated from evoked responses. Third, Remember responses were relatively few per participant and condition (Tables S1 and S2 in Text S1), which limits the precision of the trial-wise estimates.

Being the agent of an action changed how an event was encoded. Choosing what to act on improved later memory for that item, most for recollection, and more weakly for the item not chosen. With fluent control over the outcome, this recollection advantage also lasted better over a day. During encoding, choosing raised phase alignment early in the action, and later theta alignment distinguished what would be recollected from what would only feel familiar. Part of what makes a memory one’s own may therefore be set at the moment of choosing.

## Author contributions

Q.C. and J.S. designed research; Q.C. performed research and analyzed data; J.S. advised on data analysis; Q.C. and J.S. wrote the paper.

## Acknowledgments

This study was supported by the Royal Society Leverhulme Trust Senior Research Fellowship (SRF\R1\241015) to J.S.

For the purpose of open access, the author has applied a Creative Commons Attribution (CC BY) licence to any Author Accepted Manuscript version arising from this submission.

## Open Science Framework preregistration

https://doi.org/10.17605/OSF.IO/Z43DF

---

**Text S1**. Supplementary methods and results. Additional behavioral and EEG analyses, response-initiation-latency controls, Bayesian model checks, and supplementary figures and tables.

## References

Backus AR, Schoffelen J-M, Szebényi S, Hanslmayr S, Doeller CF (2016) Hippocampal-prefrontal theta oscillations support memory integration. Curr Biol 26:450–457.

Chambon V, Sidarus N, Haggard P (2014) From action intentions to action effects: how does the sense of agency come about? Front Hum Neurosci 8:320.

Clouter A, Shapiro KL, Hanslmayr S (2017) Theta phase synchronization is the glue that binds human associative memory. Curr Biol 27:3143–3148.e6.

Conway MA (2005) Memory and the self. J Mem Lang 53:594–628.

Conway MA, Dewhurst SA (1995) The self and recollective experience. Appl Cogn Psychol 9:1–19.

Conway MA, Pleydell-Pearce CW (2000) The construction of autobiographical memories in the self-memory system. Psychol Rev 107:261–288.

Conway MA, Singer JA, Tagini A (2004) The self and autobiographical memory: correspondence and coherence. Soc Cogn 22:491–529.

Cunningham SJ, Brady-Van Den Bos M, Turk DJ (2011) Exploring the effects of ownership and choice on self-memory biases. Memory 19:449–461.

Dewey JA, Pacherie E, Knoblich G (2014) The phenomenology of controlling a moving object with another person. Cognition 132:383–397.

DuBrow S, Eberts EA, Murty VP (2019) A common mechanism underlying choice’s influence on preference and memory. Psychon Bull Rev 26:1958–1966.

Elliott BL, Blais C, McClure SM, Brewer GA (2020) Neural correlates underlying the effect of reward value on recognition memory. Neuroimage 206:116296.

Estudillo AJ, Bindemann M (2014) Generalization across view in face memory and face matching. Iperception 5:589–601.

Fell J, Ludowig E, Rosburg T, Axmacher N, Elger CE (2008) Phase-locking within human mediotemporal lobe predicts memory formation. Neuroimage 43:410–419.

Gallagher S (2000) Philosophical conceptions of the self: implications for cognitive science. Trends Cogn Sci 4:14–21.

Guderian S, Düzel E (2005) Induced theta oscillations mediate large-scale synchrony with mediotemporal areas during recollection in humans. Hippocampus 15:901–912.

Guth TA, Brandt A, Reinacher PC, Schulze-Bonhage A, Jacobs J, Kunz L (2025) Theta-phase locking of single neurons during human spatial memory. Nat Commun 16:7402.

Haggard P (2017) Sense of agency in the human brain. Nat Rev Neurosci 18:196–207.

Haggard P, Chambon V (2012) Sense of agency. Curr Biol 22:R390–R392.

Herweg NA, Apitz T, Leicht G, Mulert C, Fuentemilla L, Bunzeck N (2016) Theta-alpha oscillations bind the hippocampus, prefrontal cortex, and striatum during recollection: evidence from simultaneous EEG-fMRI. J Neurosci 36:3579–3587.

Heusser AC, Poeppel D, Ezzyat Y, Davachi L (2016) Episodic sequence memory is supported by a theta–gamma phase code. Nat Neurosci 19:1374–1380.

Jutras MJ, Fries P, Buffalo EA (2013) Oscillatory activity in the monkey hippocampus during visual exploration and memory formation. Proc Natl Acad Sci U S A 110:13144–13149.

Lafleur A, Soulières I, Forgeot d’Arc B (2020) Sense of agency: sensorimotor signals and social context are differentially weighed at implicit and explicit levels. Conscious Cogn 84:103004.

Lakatos P, Gross J, Thut G (2019) A new unifying account of the roles of neuronal entrainment. Curr Biol 29:R890–R905.

Lawrence R, Chai XJ (2021) Self-referential encoding of source information in recollection memory. PLoS One 16:e0248044.

Li J, Chen S, Zhang L, Weng L, Lin X, Tu Y, Peng W (2026) Dynamic expectation strength and precision shape human pain perception through shared and dissociable α-oscillatory mechanisms. PLoS Biol 24:e3003675.

Lisman JE, Jensen O (2013) The theta-gamma neural code. Neuron 77:1002–1016.

Long Z, Northoff G, Fu X (2025) Task difficulty modulates the effect of mind wandering on phase dynamics. Proc Natl Acad Sci U S A 122:e2416387122.

Lu X, Bilkey DK (2010) The velocity-related firing property of hippocampal place cells is dependent on self-movement. Hippocampus 20:573–583.

Markant DB, Ruggeri A, Gureckis TM, Xu F (2016) Enhanced memory as a common effect of active learning. Mind Brain Educ 10:142–152.

Murty VP, DuBrow S, Davachi L (2015) The simple act of choosing influences declarative memory. J Neurosci 35:6255–6264.

Nyhus E, Curran T (2010) Functional role of gamma and theta oscillations in episodic memory. Neurosci Biobehav Rev 34:1023–1035.

Oostenveld R, Fries P, Maris E, Schoffelen J-M (2011) FieldTrip: open source software for advanced analysis of MEG, EEG, and invasive electrophysiological data. Comput Intell Neurosci 2011:156869.

Richter CG, Thompson WH, Bosman CA, Fries P (2015) A jackknife approach to quantifying single-trial correlation between covariance-based metrics undefined on a single-trial basis. Neuroimage 114:57–70.

Sato A, Yasuda A (2005) Illusion of sense of self-agency: discrepancy between the predicted and actual sensory consequences of actions modulates the sense of self-agency, but not the sense of self-ownership. Cognition 94:241–255.

Schwarz KA, Weller L, Klaffehn AL, Pfister R (2019) The effects of action choice on temporal binding, agency ratings, and their correlation. Conscious Cogn 75:102807.

Sidarus N, Chambon V, Haggard P (2013) Priming of actions increases sense of control over unexpected outcomes. Conscious Cogn 22:1403–1411.

Siebenhühner F, Wang SH, Palva JM, Palva S (2016) Cross-frequency synchronization connects networks of fast and slow oscillations during visual working memory maintenance. Elife 5:e13451.

Song Z, Wixted JT, Hopkins RO, Squire LR (2011) Impaired capacity for familiarity after hippocampal damage. Proc Natl Acad Sci U S A 108:9655–9660.

Stephens M (2017) False discovery rates: a new deal. Biostatistics 18:275–294.

Sui J, Humphreys GW (2015) The integrative self: how self-reference integrates perception and memory. Trends Cogn Sci 19:719–728.

Synofzik M, Vosgerau G, Newen A (2008) Beyond the comparator model: a multifactorial two-step account of agency. Conscious Cogn 17:219–239.

Tallon-Baudry C, Bertrand O (1999) Oscillatory gamma activity in humans and its role in object representation. Trends Cogn Sci 3:151–162.

Tort ABL, Komorowski RW, Manns JR, Kopell NJ, Eichenbaum H (2009) Theta–gamma coupling increases during the learning of item–context associations. Proc Natl Acad Sci U S A 106:20942–20947.

Tulving E (1983) Elements of episodic memory. Oxford: Oxford University Press.

Tulving E (1985) Memory and consciousness. Can Psychol 26:1–12.

Tulving E (2002) Episodic memory: from mind to brain. Annu Rev Psychol 53:1–25.

Vanderwolf CH (1969) Hippocampal electrical activity and voluntary movement in the rat. Electroencephalogr Clin Neurophysiol 26:407–418.

Voss A, Rothermund K, Gast A, Wentura D (2013) Cognitive processes in associative and categorical priming: a diffusion model analysis. J Exp Psychol Gen 142:536–559.

Voss JL, Gonsalves BD, Federmeier KD, Tranel D, Cohen NJ (2011) Hippocampal brain-network coordination during volitional exploratory behavior enhances learning. Nat Neurosci 14:115–120.

Wegner DM (2004) Précis of *The illusion of conscious will*. Behav Brain Sci 27:649–659.

Wegner DM, Wheatley T (1999) Apparent mental causation: sources of the experience of will. Am Psychol 54:480–492.

Wenke D, Fleming SM, Haggard P (2010) Subliminal priming of actions influences sense of control over effects of action. Cognition 115:26–38.

Wheeler MA, Stuss DT, Tulving E (1997) Toward a theory of episodic memory: the frontal lobes and autonoetic consciousness. Psychol Bull 121:331–354.

White CN, Poldrack RA (2014) Decomposing bias in different types of simple decisions. J Exp Psychol Learn Mem Cogn 40:385–398.

Wixted JT (2007) Dual-process theory and signal-detection theory of recognition memory. Psychol Rev 114:152–176.

Xie S, Singer J, Yilmaz B, Kaiser D, Cichy RM (2025) Recurrence affects the geometry of visual representations across the ventral visual stream in the human brain. PLoS Biol 23:e3003354.

Yonelinas AP (1994) Receiver-operating characteristics in recognition memory: evidence for a dual-process model. J Exp Psychol Learn Mem Cogn 20:1341–1354.

Yonelinas AP (2001) Components of episodic memory: the contribution of recollection and familiarity. Philos Trans R Soc Lond B Biol Sci 356:1363–1374.

Yonelinas AP (2002) The nature of recollection and familiarity: a review of 30 years of research. J Mem Lang 46:441–517.

Yonelinas AP, Jacoby LL (1995) The relation between remembering and knowing as bases for recognition: effects of size congruency. J Mem Lang 34:622–643.

Yonelinas AP, Parks CM (2007) Receiver operating characteristics (ROCs) in recognition memory: A review. Psychol Bull 133:800–832.

