## Supplemental Materials for "Self-generated action shapes theta phase alignment at encoding and later recollection"

### Text S1

#### Confidence ROC model comparison across experiments

To determine whether selected-item confidence responses were better captured by single- or dual-process accounts, the equal-variance signal-detection (EVSD), unequal-variance signal-detection (UVSD) and dual-process signal-detection (DPSD) models were compared (Yonelinas, 1994; Wixted, 2007; Yonelinas and Parks, 2007). The primary analysis pooled delayed-test data across Experiments 1–3 ($N$ = 135).

Model-selection criteria provided converging but criterion-dependent evidence. DPSD was preferred by both Akaike information criterion (AIC) and leave-one-out information criterion (LOOIC): relative to DPSD, the losses were ΔAIC = 21.89 and ΔLOOIC = 18.74 for UVSD, and ΔAIC = 41.79 and ΔLOOIC = 35.33 for EVSD. All pooled PSIS-LOO estimates had acceptable Pareto-$k$ diagnostics ($\leq$ .70). By contrast, the Bayesian information criterion (BIC) favored the more parsimonious UVSD model, with ΔBIC = 2.48 for DPSD and 11.78 for EVSD. Thus, the pooled comparison favored DPSD under criteria emphasizing fit and out-of-sample predictive performance, whereas the stronger complexity penalty imposed by BIC retained support for UVSD.

Posterior-predictive checks showed the same overall ordering. Posterior-predictive quantities were derived from a Laplace approximation. DPSD produced the lowest ROC root-mean-square error (RMSE = .008), compared with UVSD (.011) and EVSD (.012), and the highest coverage of the observed ROC coordinates by the 95% posterior-predictive intervals (75%, compared with 70% and 60%, respectively). DPSD also produced the smallest response-category likelihood-ratio discrepancy ($G^2$ = 38.93), followed by UVSD (66.82) and EVSD (88.72). Consistent with these summary measures, DPSD prediction residuals were generally closer to zero across the confidence thresholds.

Separate-analysis AIC comparisons indicated that the DPSD preference was reproducible across the delayed tests. DPSD was preferred in Experiment 1 delayed (ΔAIC: UVSD = 2.00, EVSD = 22.53), Experiment 2 delayed (UVSD = 12.09, EVSD = 12.21) and Experiment 3 delayed (UVSD = 3.83, EVSD = 7.46). The Experiment 2 immediate test was an exception: UVSD was preferred, with ΔAIC = 8.02 for DPSD and 29.94 for EVSD. Together, these results provide converging evidence that the delayed selected-item confidence ROC is better characterized by a dual-process model, while the divergent BIC result and the Experiment 2 immediate-test comparison indicate that this support is not criterion- or context-uniform.

#### Supplemental source-level analyses

##### Source-level theta and alpha phase alignment

To characterize the estimated cortical distribution of action-onset-locked phase alignment, cortical source-space cluster permutation tests of inter-trial phase coherence (ITPC) were run from 0 to 1,000 ms after action onset. Separate tests examined the main effects of Selection and Feedback and their interaction.

Source-level theta ITPC showed a positive Selection effect, with stronger phase alignment following self-generated than externally generated actions. This cluster comprised 362 unique voxels, extended from 30 to 450 ms, and peaked at 270 ms in the left lingual cortex (MNI: -15, -80, 0; $p_{\text{cluster}} < .001$). Its center of mass was located near the midline (MNI: -.5, -79.0, 7.0) (**Fig. S1A**). A smaller negative Feedback cluster indicated lower theta ITPC under disrupted than fluent feedback. This cluster comprised 24 voxels, extended from 350 to 430 ms, and peaked at 370 ms in the right inferior temporal cortex (MNI: 55, -20, -20; $p_{\text{cluster}} = .047$) (**Fig. S1B**). No theta Selection × Feedback cluster survived cluster-level correction.

Alpha ITPC likewise showed a positive Selection effect, indicating stronger phase alignment following self-generated than externally generated actions. The cluster comprised 55 voxels, extended from 70 to 290 ms, and peaked at 210 ms in the right cuneus (MNI: 5, -70, 20, $p_{\text{cluster}} < .001$). Its center of mass was located near the midline (MNI: 1.6, -74.5, 35.9) (**Fig. S1C**). Alpha ITPC also showed two negative Feedback clusters. The larger cluster comprised 219 voxels, extended from 250 to 990 ms, and peaked at 450 ms in the right calcarine cortex (MNI: 25, -100, 0, $p_{\text{cluster}} < .001$). Its spatial extent encompassed broad bilateral occipital and visual cortical regions (**Fig. S1D**). A second cluster comprised 23 unique voxels, extended from 330 to 450 ms, and peaked at 390 ms in the left insula (MNI: -35, 0, 0, $p_{\text{cluster}} = .035$). This cluster additionally involved the left Rolandic operculum and superior temporal cortex (**Fig. S1E**). Both clusters indicated lower alpha ITPC under disrupted than fluent feedback. A positive alpha Selection × Feedback interaction cluster survived cluster-level correction. This cluster comprised 24 unique voxels, extended from 170 to 310 ms, and peaked at 250 ms in the left superior occipital cortex (MNI: -15, -80, 40; $p_{\text{cluster}} = .033$). The positive interaction indicated that the disrupted-minus-fluent feedback difference was greater for self-generated than externally generated actions.

Thus, self-generated action was associated with stronger early source-level phase alignment in both frequency bands, whereas Feedback-related reductions were most extensive in the alpha band. Alpha phase alignment additionally showed a localized Selection × Feedback interaction in posterior cortical regions. These condition-level phase-alignment effects motivated the analysis of theta–gamma coupling between estimated cortical sources.

**Condition-level theta–gamma phase–amplitude coupling**

Source-level theta–gamma PAC was quantified using normalized mean vector length (MVL). Selection, Feedback, and their interaction were tested separately using cluster-based permutation tests, with BH–FDR correction across the eight theta-phase seeds within each effect family. Two Selection × Feedback interaction clusters survived this correction (**Fig. S2A–D**).

The right posterior-cingulate theta-phase seed showed a positive interaction with gamma amplitude in a distributed left frontoparietal target cluster (112 voxels; peak MNI: -25, 10, 60). This effect was detected in the 100–600-ms analysis window, centered at 350 ms ($p_{\text{cluster}} = .002$, $q_{\text{FDR}} = .007$; **Fig. S2A, C**). The disrupted-minus-fluent feedback contrast was greater for self-generated than externally generated actions.

The left calcarine theta-phase seed showed a negative interaction with gamma amplitude in a left anterior frontal target cluster (114 voxels; peak MNI: -25, 30, 10). The effect was detected in the 300–800-ms analysis window, centered at 550 ms ($p_{\text{cluster}}$ = .002, $q_{\text{FDR}}$ = .007; **Fig. S2B**, **D**). Here, the disrupted-minus-fluent feedback contrast was smaller for self-generated than externally generated actions. These intervals denote the 500-ms PAC estimation windows rather than precise effect onset and offset times.

Neither the Selection nor the Feedback main effect survived the corresponding eight-seed correction for normalized MVL. The two interaction clusters defined the seed-target combinations and time windows used to construct the subsequent trial-wise jPAC predictors. Anatomical labels describe the estimated source locations and should be interpreted in light of source-localization uncertainty.

*Raw-MVL sensitivity analysis of theta–gamma coupling.* The primary theta–gamma PAC analysis described above used normalized mean vector length (MVL). To assess whether its conclusions depended on normalization, the same seed-to-whole-brain analyses were repeated using raw MVL. Selection, Feedback, and Selection × Feedback were treated as separate effect families, with Benjamini–Hochberg false-discovery-rate correction applied across the 8 theta-phase seeds within each family.

The negative Selection × Feedback interaction originating from the left calcarine seed was also present in the raw-MVL analysis. The corresponding target cluster comprised 102 voxels, was expressed in the window from 300 to 800 ms and peaked in anterior frontal cortex (MNI: -25, 30, 10, $p_{\text{cluster}}$ = .002). The seed-level interaction test survived correction across the 8 theta seeds ($q_{\text{FDR}}$ = .014). This effect was spatially and directionally consistent with the normalized-MVL interaction described above. Raw MVL additionally showed a Selection effect originating from the literature-defined right pre-supplementary motor area seed (pre-SMA). The seed-level test survived the eight-seed correction ($q_{\text{FDR}}$ = .030) and contained two negative clusters, indicating lower raw coupling following self-generated than externally generated actions. One cluster comprised 63 voxels over the same window and peaked at MNI -5, -20, 80 ($p_{\text{cluster}}$ = .004). The second comprised 30 voxels over the same window and peaked at MNI -25, -70, -10 ($p_{\text{cluster}}$ = .017). These pre-SMA effects were not present in the primary normalized-MVL results and were therefore treated as metric-dependent. Conversely, the positive posterior-cingulate-seeded interaction identified using normalized MVL was not recovered in the raw-MVL analysis. No other raw-MVL Selection, Feedback, or Selection × Feedback test survived the corresponding within-effect eight-seed correction.

*Trial-shuffled surrogate analysis of theta–gamma coupling*. A further test asked whether the two normalized-MVL Selection × Feedback interactions described above exceeded coupling expected after disrupting the trial-wise correspondence between theta phase and gamma amplitude. Within each participant and experimental condition, gamma-amplitude trials were reassigned to different theta-phase trials using 1,000 random derangements. Group inference was based on the participant-level difference between the observed interaction and the mean surrogate interaction, with two-sided sign-flip tests and BH-FDR correction across the two targeted clusters.

For the posterior-cingulate-seed cluster, the observed interaction was larger than the surrogate interaction (observed mean = .02; surrogate mean = .01; observed-minus-surrogate difference = .01, 95% CI [.01, .02], $q_{\text{FDR}} < .001$). The negative calcarine-seeded interaction was also stronger than its surrogate counterpart (observed mean = -.02; surrogate mean = -.01; observed-minus-surrogate difference = -.01, 95% CI [-.01, -.00], $q_{\text{FDR}} < .001$). Thus, both normalized-MVL interactions retained their original directions and exceeded the effects expected after disrupting within-condition trial-wise phase–amplitude correspondence. Because these tests were restricted to clusters selected in the primary PAC analysis, they constitute post-selection sensitivity analyses rather than independent confirmations.

*Trial-wise jPAC–memory associations*. Trial-wise jPAC predictors were computed within participant × Selection × Feedback groups, pooling across subsequent memory responses. Both normalized jPAC predictors entered the joint Bayesian multinomial model alongside the eight jITPC predictors, Selection, Feedback, and their interaction. Calcarine–frontal jPAC showed uncertain associations with Know versus Forgotten ($\beta$ = .02, 95% CrI [-.04, .09]), Remember versus Forgotten ($\beta$ = .02, 95% CrI [-.09, .12]), and Remember versus Know responses ($\beta$ = -.01, 95% CrI [-.12, .10]). Posterior-cingulate–cortical jPAC likewise showed uncertain associations with Know versus Forgotten ($\beta$ = -.06, 95% CrI [-.13, .003]), Remember versus Forgotten ($\beta$ = -.08, 95% CrI [-.20, .03]), and Remember versus Know responses ($\beta$ = -.02, 95% CrI [-.13, .09]). All corresponding 95% credible intervals also included zero after adjustment for target-region gamma amplitude.

**Exploratory source-level alpha–gamma phase–amplitude coupling**

Because alpha–gamma PAC was not part of the preregistered analysis, it was examined as an exploratory frequency-specific analysis. Alpha-phase seeds were derived from the source-level alpha ITPC clusters using the same leave-one-participant-out search procedure applied in the theta analysis. Coupling between 8–13 Hz alpha phase and 30–80 Hz gamma amplitude was quantified using normalized MVL. Selection, Feedback, and their interaction were treated as separate families, and BH-FDR correction was applied across the five alpha seeds within each effect family. No additional cross-metric or global correction was used for the claims reported here.

All effects that survived the five-seed correction were negative Feedback effects, indicating lower normalized alpha–gamma coupling under disrupted than fluent feedback. The test originating from the right middle occipital seed (MNI: 25, -90, 10) showed the strongest evidence and survived the five-seed correction at $q_{\text{FDR}} < .001$. It contained a large target cluster comprising 1,149 unique voxels across windows centered from 450 to 750 ms, with a peak at MNI 5, -20, 50 ($p_{\text{cluster}} < .001$) and a smaller 15-voxel cluster across windows centered from 450 to 650 ms, peaking at MNI 5, -100, 10 ($p_{\text{cluster}} = .043$).

The left calcarine seed (MNI: -5, -70, 10) also showed a negative Feedback effect that survived the five-seed correction ($q_{\text{FDR}} = .016$). This test contained two clusters. The first comprised 31 unique voxels across windows centered from 450 to 750 ms and peaked at MNI 15, -40, 40 ($p_{\text{cluster}} = .009$). The second comprised 36 voxels across windows centered from 550 to 650 ms and peaked at MNI 35, -20, 20 ($p_{\text{cluster}} = .012$). Finally, the right precuneus seed (MNI: 15, -50, 30) showed a 33-voxel negative Feedback cluster across windows centered from 550 to 750 ms, peaking at MNI 55, 30, -10 ($p_{\text{cluster}}$ = .008). This seed-level test also survived the five-seed correction ($q_{\text{FDR}}$ = .016). No normalized-MVL Selection or Selection × Feedback test survived the corresponding five-seed correction.

*Trial-shuffled surrogate analysis of alpha–gamma coupling*. The five normalized alpha–gamma clusters were subsequently subjected to a targeted surrogate sensitivity analysis. For clusters spanning multiple overlapping PAC windows, the spatial masks at the cluster's peak window were fixed before computing the surrogate statistics. Within each participant and condition, 1,000 trial derangements disrupted the trial-wise correspondence between alpha phase and gamma amplitude. Group inference used two-sided sign-flip tests to assess whether the observed disrupted-minus-fluent feedback contrasts differed from the corresponding surrogate contrast, with BH–FDR correction across the five frozen targets. Mean differences and 95% t-based confidence intervals were calculated across 42 participants.

Four of the five Feedback effects remained more negative than their corresponding surrogate effects after correction. For the first calcarine-seeded cluster, the observed Feedback effect was -.014 compared with a surrogate mean of -.01 (observed-minus-surrogate difference = -.0069, 95% CI [-.012, -.00], $q_{\text{FDR}}$ = .010). The second calcarine-seeded cluster showed a comparable result (observed = -.01; surrogate = -.01; difference = -.01, 95% CI [-.01, -.00], $q_{\text{FDR}}$ = .010).

The large middle-occipital-seeded cluster also retained a reliable surrogate-corrected Feedback effect (observed = -.02; surrogate = -.01; difference = -.00, 95% CI [-.01, -.00], $q_{\text{FDR}}$ = .005). In contrast, its smaller posterior target cluster did not differ reliably from the surrogate contrast (observed = -.01; surrogate = -.01; difference = -.00, 95% CI [-.01, .00], $q_{\text{FDR}}$ = .501). Finally, the precuneus-seeded cluster remained reliably more negative than its surrogate counterpart (observed = -.02; surrogate = -.01; difference = -.01, 95% CI [-.01, .00], $q_{\text{FDR}}$ = .002).

Thus, four of the five exploratory alpha–gamma Feedback clusters exceeded the effect expected after disrupting trial-wise alpha-phase/gamma-amplitude correspondence. The exception was the smaller posterior cluster associated with the middle-occipital seed. These results provide a sensitivity check on the detected alpha–gamma effects but, because both the alpha–gamma analysis and the targeted surrogate tests were exploratory and the surrogate targets were selected from the original results, they should not be interpreted as independent confirmation.

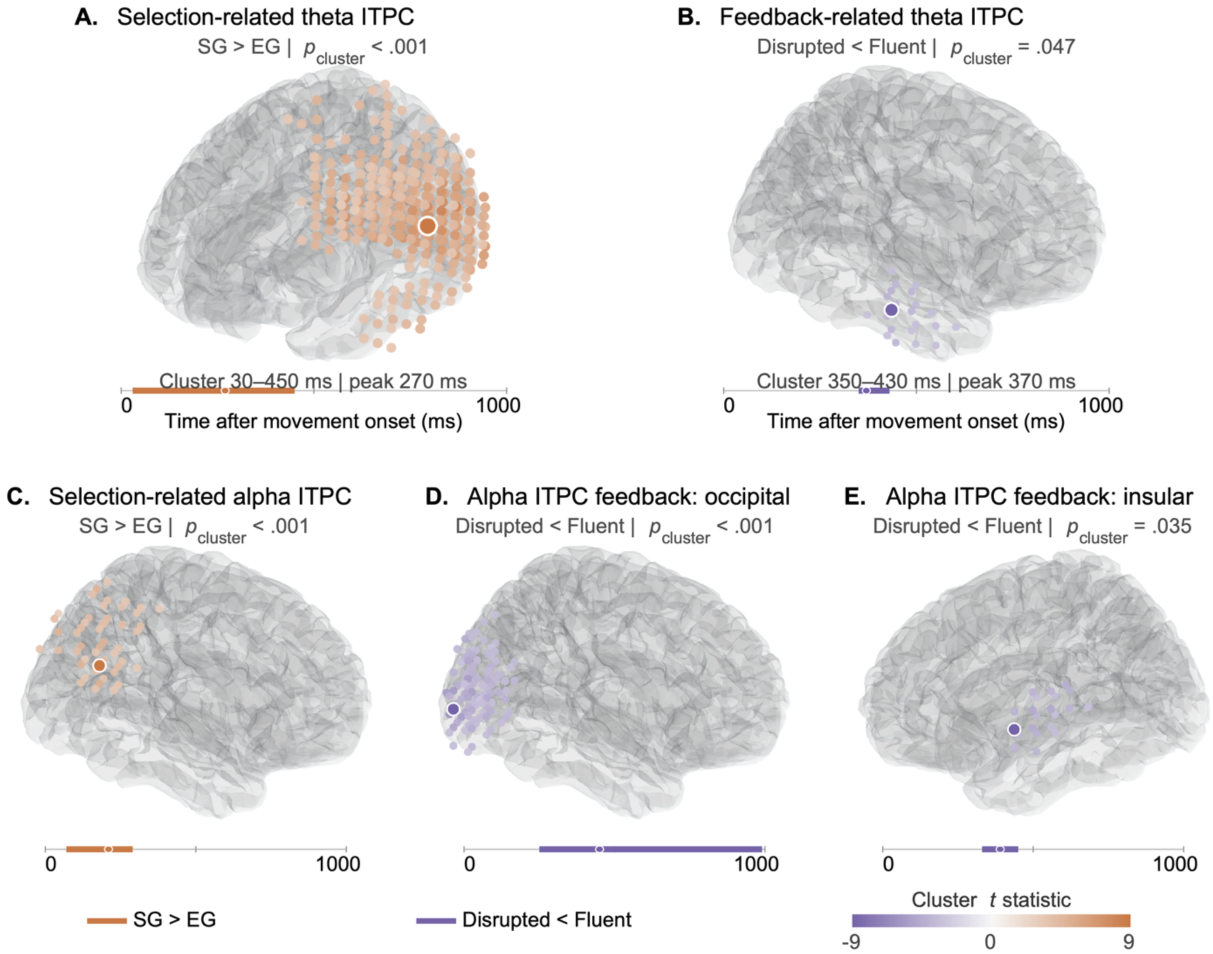


**Figure S1. Self-generated action enhances, whereas disrupted feedback reduces, source-level theta and alpha phase alignment.**

Related to **Fig. 3**. Source-level inter-trial phase coherence (ITPC) during the action-onset-locked interval from 0 to 1,000 ms. **A,** Theta ITPC was greater for self-generated than externally generated actions from 30–450 ms (peak, 270 ms; $p_{\text{cluster}} < .001$). **B**, Theta ITPC was lower under disrupted than fluent feedback from 350–430 ms (peak, 370 ms; $p_{\text{cluster}} = .047$). **C**, Alpha ITPC was greater for self-generated than externally generated actions from 70–290 ms (peak, 210 ms; $p_{\text{cluster}} < .001$). **D**, Alpha ITPC was lower under disrupted than fluent feedback in a broad bilateral occipital/visual cluster (250–990 ms; peak, 450 ms; $p_{\text{cluster}} < .001$). **E,** A second reduction was observed in a left insular–opercular cluster (330–450 ms; peak, 390 ms; $p_{\text{cluster}} = .035$). Surface renderings show voxels within an 80-ms window centered on each cluster peak. The larger, white-edged colored circle marks the peak voxel. **A** and **E** are shown from left-lateral perspectives, whereas **B–D** are shown from right-lateral perspectives. The right-lateral views in **C** and **D** highlight their right-sided peaks, although both clusters extend bilaterally. Voxel colors represent the signed voxel-wise *t* statistic averaged across the displayed 80-ms window, using a common diverging scale shared across all five panels. Orange denotes positive *t* statistics and purple denotes negative *t* statistics, corresponding here to self-generated > externally generated and disrupted feedback < fluent feedback effects, respectively. Greater color saturation indicates a larger absolute t statistic. Colored horizontal bars show the temporal extent of each cluster, and the small, white-edged circle marks the cluster peak time. All displayed clusters survived whole-cortical-source-space spatiotemporal cluster-based permutation correction.

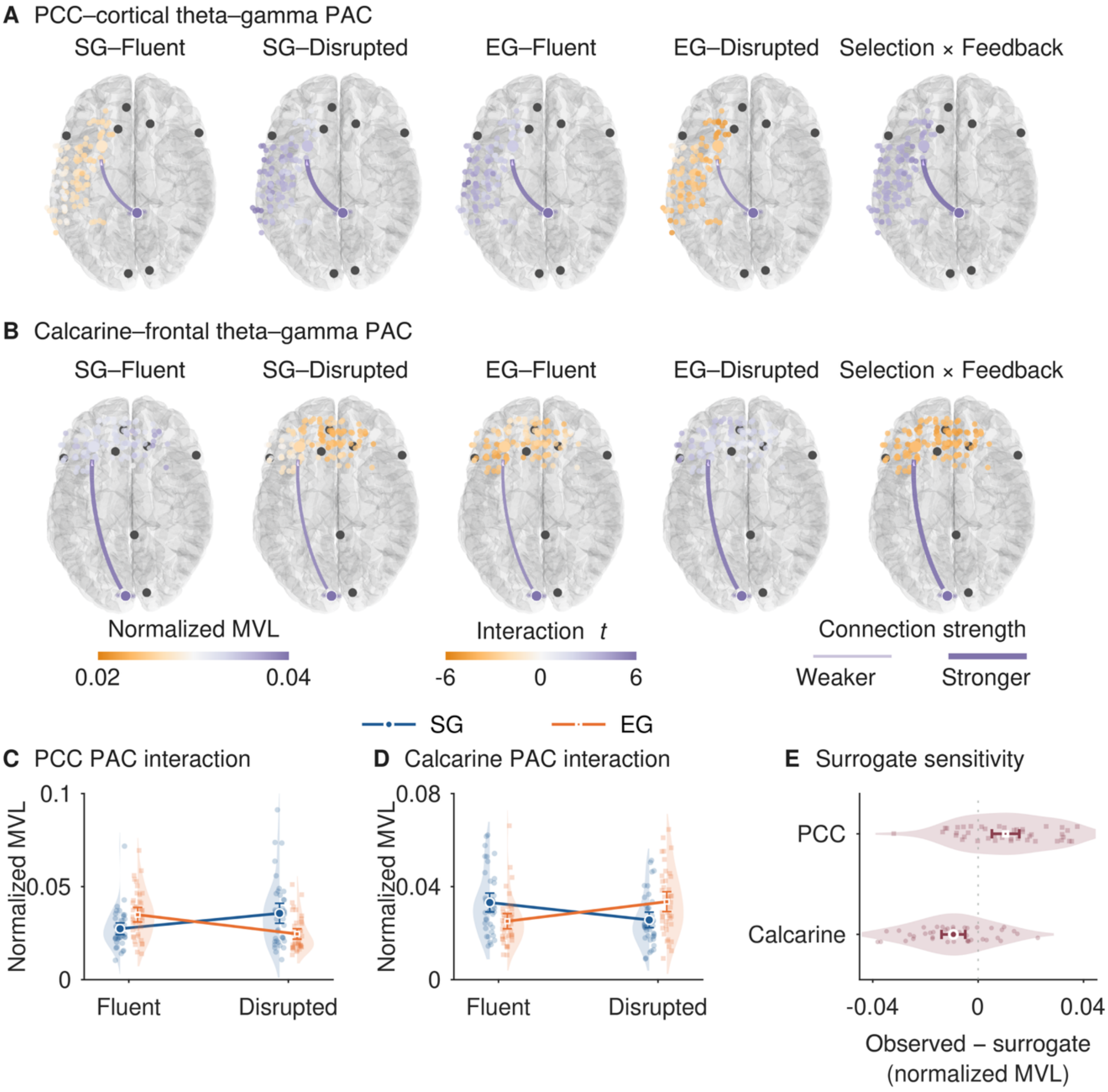


**Figure S2. Source-level theta–gamma phase–amplitude coupling.**

**A** and **B**, Dorsal source-space maps of theta–gamma phase–amplitude coupling (PAC) for posterior-cingulate–cortical (**A**) and calcarine–frontal (**B**) networks. The first four maps show normalized mean vector length for SG–Fluent, SG–Disrupted, EG–Fluent, and EG–Disrupted conditions, and the fifth map shows the Selection × Feedback interaction. Connection width and opacity encode coupling strength. **C** and **D**, Participant-level normalized coupling values underlying the corresponding interactions. Small points show participants, half violins show distributions and connected symbols and error bars show group means and 95% confidence intervals. **E**, Targeted surrogate sensitivity analysis showing participant-level observed-minus-surrogate interaction differences. Small points show participants, half-violins show distributions, and large symbols and intervals indicate means and 95% confidence intervals. n = 42 participants.

#### Response-initiation latency and latency-adjusted trial-wise neural-memory models in Experiment 3

Response-initiation latency was defined as the interval from stimulus onset to the response-start marker preceding movement. Condition summaries were based on participant-level medians across 6,679 retained EEG epochs from 42 participants. Median latency was approximately 1.02–1.03 s for self-generated trials and .70–.74 s for externally generated trials. Movement duration was obtained from the behavioral task records (`ActualTimeMoving`) and remained tightly constrained at approximately 2.06 s across the four experimental conditions.

*Latency-matched condition-level ITPC sensitivity analysis*. Because conventional ITPC is estimated across trials and therefore cannot accommodate a trial-level covariate directly, a response-initiation-latency-matched sensitivity analysis was run instead. Within each participant and Feedback condition, self-generated and externally generated trials were matched on log-transformed response-initiation latency using a pooled-stratum standardized caliper of .20 SD. Participants were retained when at least 15 matched pairs were available in both Feedback conditions. The criterion retained 35 of the 42 participants and 1,839 matched trial pairs. Matching produced close latency balance: across the 70 participant-by-Feedback strata in the retained sample, the absolute difference between self-generated and externally generated trials in mean standardized log latency had a median of .02, a 90$^{th}$ percentile of .05 and a maximum of .70. The corresponding absolute difference in mean latency had a median of 6.58 ms, a 90$^{th}$ percentile of 16.80 ms and a maximum of 28.66 ms.

Action-onset-locked ITPC was then recomputed separately for the four Selection × Feedback conditions using the time-frequency parameters of the primary analysis. Disrupted- and fluent-feedback estimates were combined using matched-pair-count weights that were identical for self-generated and externally generated trials, and the contrasts between self-generated and externally generated trials were evaluated using whole-scalp channel × time cluster-based permutation tests with 10,000 randomizations. The positive theta Selection effect remained significant after latency matching, extending from 48 to 373 ms across 42 electrodes and peaking at POz at 250.98 ms ($t$ = 8.27, two-sided $p_{\text{cluster}}$ < .001, $q_{\text{FDR}}$ < .001). The positive alpha Selection effect was likewise retained from 134 to 209 ms across 11 electrodes, with peak at PO3 at 146 ms ($t$ = 5.13, two-sided $p_{\text{cluster}}$ < .001, $q_{\text{FDR}}$ < .001).

As a targeted sensitivity analysis, the latency-matched differences between self-generated and externally generated ITPC were also averaged within the spatiotemporal masks of the original significant Selection clusters. The matched difference remained positive within the original theta mask (19–416 ms; 59 electrodes; $M$ = .08, 95% CI [.06, .10], $t$ (34) = 8.16, $q_{\text{FDR}}$ < .001, $d_z$ = 1.38) and the original alpha mask (127–245 ms; 39 electrodes; $M$ = .06, 95% CI [.03, .08], $t$ (34) = 6.30, $q_{\text{FDR}}$ < .001, $d_z$ = 1.06). Thus, the principal selection-related increases in action-onset-locked theta and alpha ITPC remained reliable when self-generated and externally generated trials were closely matched on response-initiation latency. Because these latter tests used masks defined from the original results, they were interpreted as post-selection sensitivity analyses rather than independent confirmations.

*Latency-adjusted trial-wise neural-memory models in Experiment 3*. A sensitivity analysis examined whether response-initiation latency accounted for the trial-wise neural–memory associations in Experiment 3. The analysis used the same 4,994 trials from 42 participants as the primary models. Neural predictors were computed within participant × Selection × Feedback groups without using subsequent memory labels. Log-transformed latency was decomposed into within-participant and between-participant components, which were fitted separately for selected-item memory and unselected-item recognition, yielding 16 latency-adjusted models. Priors, random-effects structures, and sampling settings matched the corresponding primary models. In the condition-only model adjusted for latency, self-generated action remained associated with Remember versus Forgotten responses ($\beta$ = .55, 95% CrI [.35, .77]) and Remember versus Know responses ($\beta$ = .35, 95% CrI [.13, .59]). The adjusted Selection effect on unselected-item recognition remained uncertain ($\beta$ = .04, 95% CrI [-.07, .15]).

In the joint jITPC–jPAC model, early middle-frontal alpha jITPC remained positively associated with Remember versus Forgotten responses ($\beta$ = .15, 95% CrI [.02, .28], $q_{\mathrm{BFSR}}$ = .014). Later middle-posterior theta jITPC remained positively associated with Remember versus Know responses ($\beta$ = .14, 95% CrI [.01, .27], $q_{\mathrm{BFSR}}$ = .013). Its association with Remember versus Forgotten responses remained uncertain ($\beta$ = .08, 95% CrI [-.04, .21]). Neither jPAC predictor clearly distinguished Remember from Forgotten or Know responses. Within-participant latency was positively associated with Know versus Forgotten and Remember versus Forgotten responses, but its Remember-versus-Know association remained uncertain. For unselected-item recognition, within-participant latency was positively associated with correct recognition ($\beta$ = .10, 95% CrI [.03, .17]). All ten neural predictors had 95% CrIs including zero in the joint unselected-item model. All 16 latency-adjusted models completed without divergent transitions or maximum-treedepth exceedances (maximum $\hat{R}$ = 1.009; minimum effective sample size = 756).

#### Latency-adjusted behavioral memory models in Experiments 1 and 2

Response-initiation latency was calculated from image onset in the original encoding records. Log-transformed latency was decomposed into standardized within-participant and between-participant components. Both components were added to the corresponding primary selected-item and unselected-item models, preserving their analysis samples, experimental terms, priors, and random-effects structures. Experiment 2 models retained Delay and its interactions with Selection and Feedback.

In Experiment 1, the Selection effects remained positive after latency adjustment for Know versus Forgotten ($\beta$ = .21, 95% CrI [.10, .33]), Remember versus Forgotten ($\beta$ = .45, 95% CrI [.24, .66]), and Remember versus Know responses ($\beta$ = .24, 95% CrI [.006, .46]). The Selection benefit also remained for unselected-item recognition ($\beta$ = .23, 95% CrI [.10, .38]). In Experiment 2, latency-adjusted Selection effects, averaged across Feedback, remained positive for Know versus Forgotten and Remember versus Forgotten responses at both tests. Immediate-test estimates were $\beta$ = .21, 95% CrI [.09, .33], and $\beta$ = .40, 95% CrI [.21, .59], respectively. Delayed-test estimates were $\beta$ = .20, 95% CrI [.08, .32], and $\beta$ = .38, 95% CrI [.17, .60]. The corresponding Remember-versus-Know effects were uncertain at both tests. Unselected-item recognition retained a Selection benefit immediately ($\beta$ = .19, 95% CrI [.05, .33]), but not clearly after delay ($\beta$ = .03, 95% CrI [-.11, .16]). The negative Selection × Delay interaction remained credible ($\beta$ = -.16, 95% CrI [-.26, -.06]).

#### Theta–alpha cross-frequency phase synchronization

Because both theta- and alpha-band phase consistency varied with the experimental factors, a preregistered analysis tested whether these effects were accompanied by theta–alpha cross-frequency phase synchronization. Within each channel, 2:1 theta–alpha phase-phase coupling was quantified as the consistency of the phase difference, using 300-ms sliding windows advanced in 50-ms steps across the 0–1-s interval after action onset. Theta and alpha phases were extracted at 4–7.99 Hz and 8–12.99 Hz, respectively. To control for coupling attributable to the temporal structure of the individual frequency bands, 1,000 surrogate estimates were generated by deranging alpha trials within each condition while preserving the condition-specific theta and alpha signals. Surrogate-corrected coupling was calculated as the observed coupling minus the mean surrogate coupling. Selection, Feedback, and Selection × Feedback effects were evaluated using whole-scalp channel-by-time cluster-based permutation tests with 10,000 permutations.

Surrogate-corrected theta–alpha coupling revealed a significant negative cluster for the main effect of Selection from 149 to 398 ms, involving frontocentral, centroparietal, and posterior electrodes (F5, FC3, CP3, P3, AF4, FC6, CP4, P2, P4, and PO4; $p_{cluster} = .044$). Within the identified cluster, surrogate-corrected coupling was descriptively lower following self-generated actions ($M$ = -.01, 95% CI [-.01, -.00]) than following externally generated actions ($M$ = .01, 95% CI [.00, .01]), corresponding to a self-generated-minus-externally generated difference of $\Delta R_{2:1}$ = -.01, 95% CI [-.01, -.01]. A later positive Selection cluster from 697 to 847 ms did not meet the cluster-corrected threshold ($p_{cluster} = .055$). No significant clusters were identified for Feedback or the Selection × Feedback interaction ($p_{cluster} \geq .181$).

Sensitivity analyses expressed relative to the surrogate distribution reproduced the early negative Selection cluster ($p_{cluster} = .033$) and additionally identified a later positive Selection cluster from 647 to 847 ms ($p_{cluster} = .041$). Analyses of raw, uncorrected coupling produced a feedback-related cluster ($p_{cluster} < .001$) and a Selection × Feedback interaction cluster ($p_{cluster} = .048$). However, neither effect remained after surrogate correction. These raw effects may therefore reflect condition-related differences in the temporal structure or phase consistency of the component rhythms rather than theta–alpha phase synchronization exceeding that expected from the individual frequency bands. Overall, the surrogate-corrected results indicated weaker early theta–alpha phase synchronization following self-generated than externally generated actions, without clear evidence that this coupling varied with Feedback.

#### Bayesian mixed-effects analyses for Experiment 2

To give a complete account of the Experiment 2 behavioral results, the full Bayesian mixed-effects analyses of selected-item and unselected-item recognition are reported here. Selected-item recognition was modeled as a multinomial response, with Forgotten as the reference category, allowing separate tests of Know responses, Remember responses, and the direct Remember-versus-Know comparison. Unselected-item recognition was analyzed with a

Bernoulli mixed-effects model. In both analyses, Selection (externally generated = -1; self-generated = +1), Feedback (fluent = −1, disrupted = +1), Delay (immediate = 0, delayed = 1) and their interactions were entered as fixed effects. Positive Selection coefficients indicate higher responding after self-generated than externally generated action, positive Feedback coefficients indicate higher responding after disrupted than fluent feedback, and Delay effects compare the delayed with the immediate test. Posterior means and 95% credible intervals are reported, and interactions whose 95% CrI excluded zero were followed up with posterior predicted probability contrasts.

Experiment 2 replicated the behavioral pattern observed in Experiment 1. In the combined immediate/delayed model, the Selection coefficients represented effects at the immediate test, averaged across feedback conditions. At this test, self-generated action increased Know responses relative to Forgotten responses, $\beta = .24$, 95% CrI [.12, .36]. Self-generated action also increased Remember responses relative to Forgotten responses, $\beta = .48$, 95% CrI [.30, .67]. The Selection effect was also evident in the direct Remember-versus-Know comparison, $\beta = .24$, 95% CrI [.03, .45]. Thus, self-generated action shifted recognition responses away from forgetting toward both familiarity and recollection, with the strongest shift toward recollection.

Recognition performance declined after the 24-hour delay. Compared with the immediate test, the delayed test reduced Know responses relative to Forgotten responses, $\beta = -.22$, 95% CrI [-.32, -.11], and produced a larger reduction in Remember responses relative to Forgotten responses, $\beta = -1.20$, 95% CrI [-1.38, -1.03]. The delay effect was also credible for the Remember-versus-Know comparison, $\beta = -.99$, 95% CrI [-1.17, -.81], indicating that recollection declined more sharply than familiarity over time.

The only interaction whose 95% CrI excluded zero in the categorical memory model on the selected items was the Selection × Feedback × Delay interaction for the Remember–versus–Know comparison, $\beta = -.25$, 95% CrI [-.44, -.07]. Follow-up contrasts of posterior predicted probabilities located this pattern mainly in the delayed test under disrupted feedback. After the 24-hour delay, self-generated action did not credibly change the probability of Remember responses under disrupted feedback, $\Delta M = .00$, 95% CrI [-.03, .02], but it increased the probability of Know responses, $\Delta M = .12$, 95% CrI [.04, .20]. Consequently, the Remember-Know probability difference was lower for self-generated than externally generated actions in the delayed disrupted-feedback condition, $\Delta M = -.12$, 95% CrI [-.21, -.03]. In contrast, the corresponding delayed fluent-feedback contrast was uncertain, $\Delta$ M = -.02, 95% CrI [-.11, .08]. Although the fluent-versus-disrupted difference in this follow-up contrast was in the same direction, its credible interval overlapped zero, $\Delta M = .10$, 95% CrI [-.02, .23]. No other interaction terms in the selected-item memory model were credibly different from zero.

Unselected-item recognition also showed a self-generated action benefit. Self-generated action increased unselected-item recognition relative to externally generated action, $\beta = .22$, 95% CrI [.09, .36]. Unselected-item recognition was lower after the 24-hour delay than in the immediate test, β = -.24, 95% CrI [-.35, -.14]. This Selection benefit was moderated by delay, $\beta = -.15$, 95% CrI [-.26, -.05], indicating that the self-generated advantage in unselected-item recognition was attenuated after 24 hours. Follow-up contrasts of posterior-predicted probabilities showed that self-generated action increased unselected-item recognition at the immediate test under fluent feedback, $\Delta M = .12$, 95% CrI [.03, .21]. The corresponding effect under disrupted feedback was positive, although its 95% CrI reached zero after rounding, $\Delta M = .09$, 95% CrI [.00, .19]. In contrast, the corresponding selection effects were smaller and uncertain after the 24-hour delay, both under fluent-feedback, $\Delta M = .05$, 95% CrI [-.04, .13], and under disrupted feedback, $\Delta M = .02$, 95% CrI [-.07, .11]. Thus, self-generated action benefited unselected-item recognition memory most clearly at the immediate test, whereas this benefit was reduced after delay.

#### Prior predictive checks for primary Bayesian models

Prior predictive checks evaluated whether the specified priors generated plausible outcome distributions. Checks were performed for the three behavioral *d′* outcomes (Remember, IRK-corrected Know and unselected-item *d′*) in each of the three experiments, yielding 9 behavioral models in total. Each prior-only model retained the data structure, model formula, Student–t likelihood and prior specification of its corresponding confirmatory model. Four chains were run for 6,000 iterations, including 3,000 warm-up iterations, and 2,000 prior-predictive datasets were sampled from each model. The means and standard deviations of the replicated datasets were compared with the corresponding observed summaries, both overall and across the experimental conditions.

Equivalent checks were conducted for all 16 primary confirmatory EEG models, comprising eight categorical selected-item memory models and eight Bernoulli unselected-item recognition models. The categorical models, which distinguished Forgotten, Know, and Remember responses, used an LKJ (2) prior for random-effect correlations. Four chains were run for 4,000 iterations, including 2,000 warm-up iterations, and 2,000 prior-predictive datasets were sampled from each EEG model. Prior expected probabilities and fully replicated response probabilities were compared with the observed outcome proportions, overall and separately across the Selection × Feedback conditions.

Across the behavioral models, the observed mean *d′* values ranged from .37 to .99 and fell within the model-specific 95% prior-predictive intervals. Their percentile positions within the corresponding prior-predictive distributions ranged from approximately 64.0% to 80.4%, indicating that the priors adequately covered the observed central tendencies. The observed standard deviations ranged from .33 to .53 and generally occupied the lower tail of the prior-predictive distributions (percentile range, 0.1–4.7%, with 6 of 9 models below the 2.5$^{th}$ percentile). However, all 48 condition-specific observed means and standard deviations fell within their corresponding 95% prior-predictive intervals. Thus, the priors allowed substantially greater dispersion, including occasional extreme *d′* values, than was present in the observed data. These priors were retained because they were intended to provide conservative regularization without tightly constraining the outcome variability, and the corresponding posterior models showed satisfactory sampling diagnostics. The behavioral checks therefore indicated adequate prior coverage of the observed locations, together with conservative overall prior-predictive dispersion.

For the 16 EEG models, all 160 overall and condition-specific observed probabilities fell within their corresponding 95% prior expected-probability intervals. All 160 were also covered by intervals derived from fully replicated outcomes. Prior-only EEG fits showed satisfactory sampling diagnostics, with no divergent transitions or maximum-treedepth exceedances and a maximum $\hat{R}$ of 1.003.

#### Posterior predictive checks for primary Bayesian models

Posterior predictive checks were conducted for nine behavioral *d′*models and 16 EEG models to assess whether the fitted models reproduced key features of the observed data. For each model, 2,000 posterior-predictive datasets were generated. For the nine behavioral models, the observed and replicated means, standard deviations, 2.5$^{th}$ and 97.5$^{th}$ percentiles, minima and maxima were compared, both overall and within experimental conditions. For the EEG models, observed and replicated outcome probabilities were compared overall and within the four Selection × Feedback conditions.

The behavioral models reproduced the observed central tendencies and dispersion well. All 57 observed means and 56 of the 57 observed standard deviations fell within their corresponding 95% posterior-predictive intervals. In particular, the overall mean and standard deviation were covered for all nine behavioral outcomes, including the Experiment 2 and Experiment 3 outcomes whose observed dispersion occupied the lower tail of the prior predictive distributions. Across all behavioral distributional summaries, 331 of 342 (96.8%) fell within the corresponding 95% posterior-predictive intervals. Ten of the 11 discrepancies involved minima, maxima or tail quantiles, which are especially sensitive to individual extreme observations. The remaining discrepancy was the condition-specific standard deviation of IRK-corrected Know *d′* in the Experiment 3 self-generated–fluent condition (observed = .50, 95% posterior-predictive interval [.30, .48]).

The EEG models reproduced the observed outcome proportions. Across eight selected-item categorical models and eight unselected-item Bernoulli models, all 320 observed probability summaries fell within their corresponding 95% posterior-predictive intervals. Primary EEG fits showed no divergent transitions or maximum-treedepth exceedances, with a maximum $\hat{R}$ of 1.007 and a minimum effective sample size of 551.

Although several observed behavioral standard deviations occupied the lower tail of the prior-predictive distributions, this did not translate into systematic posterior predictive misfit: the fitted models reproduced the overall mean and standard deviation of every behavioral outcome, and 56 of the 57 overall and condition-specific standard deviations fell within their corresponding 95% posterior-predictive intervals. Moreover, the substantive posterior conclusions remained stable under the weaker and more regularizing prior schemes. The prespecified primary priors were therefore retained.

#### Prior-sensitivity analysis for primary Bayesian models

To assess whether the posterior conclusions depended on the prior specification, these 25 models were refitted under two prespecified alternative schemes, yielding 50 sensitivity fits. For the behavioral models, the weaker scheme used normal (0, 1) priors for slopes, Student-t (3, 0, 2) priors for intercepts, and exponential (1) priors for scale parameters. The more regularizing scheme used normal (0, .25), Student-t (3, 0, .75), and exponential (4) priors, respectively. The gamma (2, .1) prior for the degrees-of-freedom parameter was unchanged. For the EEG models, the corresponding weaker priors were normal (0, 2), Student-t (3, 0, 5), exponential (.5), and LKJ (1). The regularizing priors were normal (0, .5), Student-t (3, 0, 1.5), exponential (2), and LKJ (4). For the categorical models, coefficient, intercept, and scale priors were applied separately to Know and Remember logits. All sensitivity fits used four chains. Behavioral models used 2,000 iterations per chain, including 1,000 warm-up iterations. EEG sensitivity fits used either 2,000 iterations with 1,000 warm-up iterations or 8,000 iterations with 4,000 warm-up iterations. Extended sampling was used for eight initial EEG sensitivity fits requiring improved convergence and for all 12 additional gamma-amplitude-control sensitivity fits. Robustness was assessed by comparing posterior means, effect directions, 95% credible intervals, sampling diagnostics, and PSIS-LOO model rankings.

Across the behavioral and updated EEG models, 252 of 255 non-intercept fixed-effect coefficients (98.8%) retained their direction across the primary and alternative prior schemes. The interval-based conclusion remained unchanged for 244 coefficients (95.7%). In the EEG models, interval-classification changes involved Feedback coefficients for Know versus Forgotten responses. EEG sensitivity fits showed no divergent transitions or maximum-treedepth exceedances, with a maximum $\hat{R}$ of 1.009. The Selection × Feedback-only model ranked first for both selected-item memory and unselected-item recognition under both alternative prior schemes. These rankings describe relative predictive performance and do not establish reliable differences between closely competing models.

#### Condition-dependent HDDM results

Hierarchical drift-diffusion modeling (HDDM) was also used to examine whether Selection and Feedback modulated the latent processes underlying Remember, Know, and unselected-item decisions. Remember and Know models contrasted selected old items receiving the corresponding response with missed old items, whereas unselected-item models contrasted correctly recognized unselected old items with missed unselected old items (**Fig. S3A**). Eight condition-dependent candidate models (M1–M8) were fitted for each outcome. M1 allowed drift rate ($v$) to vary across conditions. M2–M4 additionally allowed non-decision time ($t$), starting point ($z$) or boundary separation ($a$) to vary, respectively. M5–M7 additionally varied the pairs $t$ and $z$, $t$ and $a$, or $z$ and $a$, respectively. M8 allowed all four parameters to vary across conditions. Models were compared using the deviance information criterion (DIC), with lower values indicating better relative fit.

Among M1–M8, M1 provided the lowest DIC for Remember, Know and unselected-item models in Experiment 1 (DIC = 6,330.64, 12,257.07, and 13,901.57, respectively). M2 provided the lowest DIC for Remember, Know, and unselected-item models in Experiment 2 (DIC = 12,282.06, 23,417.68, and 26,499.75, respectively). In Experiment 3, M1 provided the lowest DIC for Remember, Know, and unselected-item models (DIC = 6,620.13, 13,015.28, and 14,733.33, respectively).

In **Experiment 1** (**Fig. S3B**), M1 indicated a higher drift rate for items following self-generated rather than externally generated action for Remember responses (*M* = .29, 95% CrI [.16, .41]) and Know responses (*M* = .19, 95% CrI [.10, .29]). Disrupted feedback was associated with a higher drift rate for Know responses (*M* = .09, 95% CrI [.0004, .19]), whereas the corresponding effect for Remember responses was uncertain (*M* = .04, 95% CrI [-.09, .17]). Neither Selection × Feedback interaction was credible (Remember: *M* = -.07, 95% CrI [-.32, .19]; Know: *M* = .02, 95% CrI [-.17, .21]). Self-generated action also increased drift rate for the corresponding unselected items (*M* = .19, 95% CrI [.10, .28]), whereas neither the Feedback effect (*M* = .05, 95% CrI [-.04, .14]) nor the Selection × Feedback interaction (*M* = -.08, 95% CrI [-.26, .10]) was credible.

In **Experiment 2** (**Fig. S3C**), the M2 summaries expressed delayed-test drift rate relative to immediate-test drift rate ($\nu_{delayed}/\nu_{immediate}$). For Remember responses, the self-generated–externally generated contrast in this ratio was not credible overall (*M* = -.74, 95% CrI [-2.98, 12.70]). Neither the Feedback effect (*M* = 2.41, 95% CrI [-10.45, 6.59]) nor the Selection × Feedback interaction (*M* = 4.99, 95% CrI [-20.81, 13.59]) was credible. The self-generated–externally generated contrast was credible only under disrupted feedback (*M* = 1.75, 95% CrI [.11, 5.13]). No main, interaction, or simple contrast was credible for Know responses. For unselected items, posterior contrasts from M2 showed no credible effects of Selection, Feedback, or their interaction with all 95% CrIs including zero.

In **Experiment 3** (**Fig.f S3D**), M1 again indicated higher drift rates following self-generated than externally generated action for Remember responses (*M* = .19, 95% CrI [.01, .36]) and Know responses (*M* = .16, 95% CrI [.05, .27]). Neither Feedback nor the Selection × Feedback interaction was credible for either response type (all 95% CrIs including zero). The self-generated–externally generated contrast on Know drift rate was credible under fluent feedback (*M* = .18, 95% CrI [.02, .33]) but not under disrupted feedback (*M* = .14, 95% CrI [-.02, .31]), whereas the corresponding Remember simple effects were not individually credible. No credible Selection, Feedback, or their interaction effect was observed for unselected items. Across the condition-dependent models, self-generated action was therefore associated with faster evidence accumulation for selected Remember and Know responses in Experiments 1 and 3, whereas delay-related effects in Experiment 2 and effects on unselected items were less consistent.

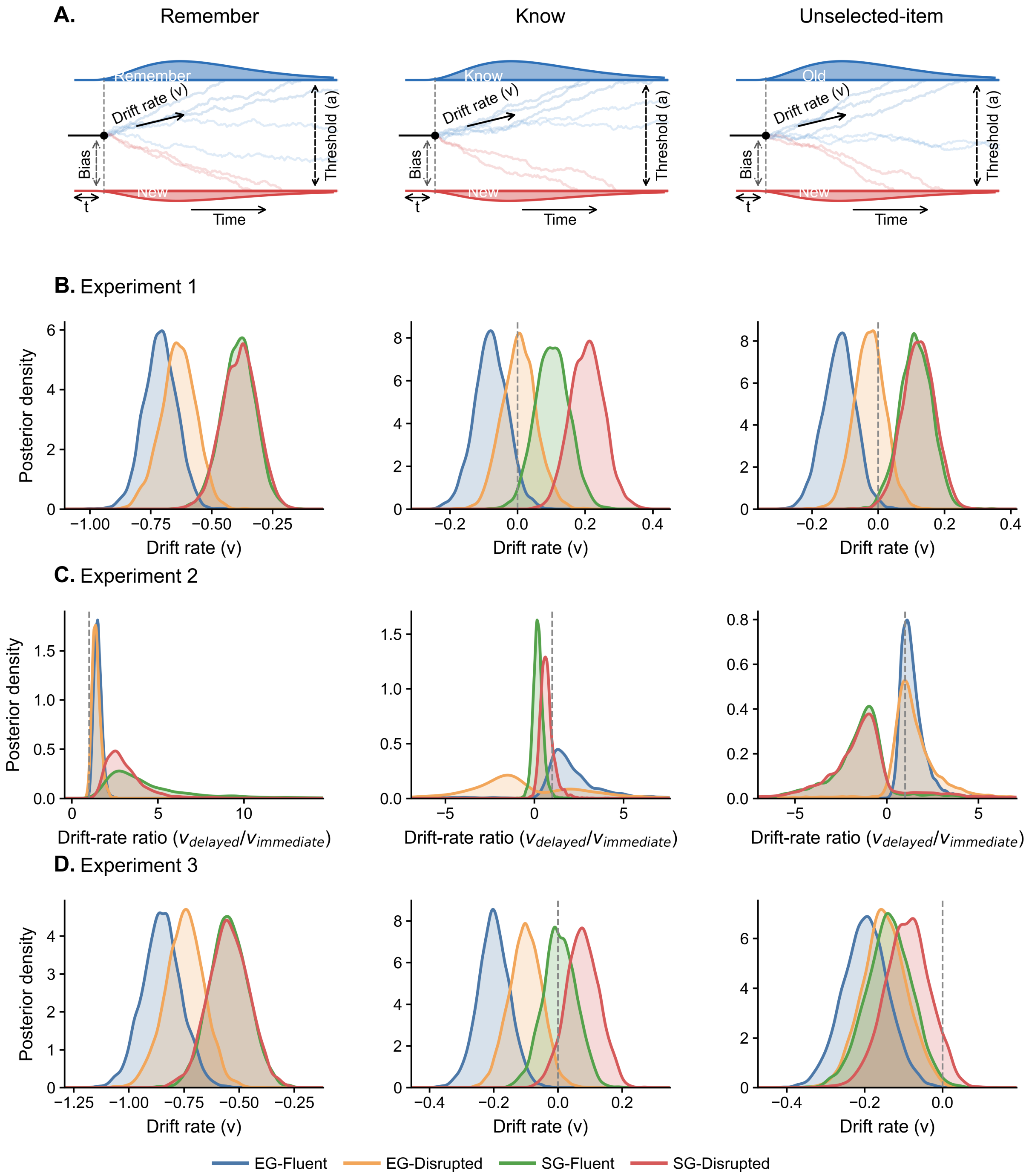


**Figure S3. HDDM schematics and posterior drift-rate distributions across experiments.**

**A**, Schematic representation of the hierarchical drift-diffusion models fitted to Remember, Know, and unselected-item decisions. The upper and lower boundaries represent the corresponding old-item response (Remember, Know, or Old) and the New response, respectively. The schematic illustrates drift rate ($v$), boundary separation ($a$), starting point ($z$), and non-decision time ($t$) and is not drawn to scale. **B**, Posterior distributions of drift rate ($v$) in Experiment 1. **C**, Posterior distributions of the delayed-to-immediate drift-rate ratio ($v_{delayed}/v_{immediate}$) in Experiment 2. **D**, Posterior distributions of drift rate ($v$) in Experiment 3. Blue, orange, green, and red curves denote externally generated (EG)–fluent, externally generated (EG)–disrupted, self-generated (SG)–fluent, and self-generated (SG)–disrupted conditions, respectively. Gray dashed lines indicate zero drift rate ($v$ = 0) in Experiments 1 and 3 and an equivalent delayed and immediate drift rate ($v_{delayed}/v_{immediate}$=1) in Experiment 2. Posterior distributions were obtained from M1 for Experiments 1 and 3 and from M2 for Experiment 2. Models were compared within the set of condition-dependent models M1–M8 using the deviance information criterion.

**Table S1. Retained action-onset-locked EEG trial counts by experimental condition and recognition response in Experiment 3**

*Related to Methods.*

| Subject ID | Age (years) | Gender | Self-generated, Disrupted feedback | | | Self-generated, Fluent feedback | | | Externally generated, Disrupted feedback | | | Externally generated, Fluent feedback | | |
|---|---|---|---|---|---|---|---|---|---|---|---|---|---|---|
| | | | Remember | Know | Forgotten | Remember | Know | Forgotten | Remember | Know | Forgotten | Remember | Know | Forgotten |
| sub05 | 21 | female | 5 | 9 | 16 | 5 | 13 | 12 | 2 | 13 | 15 | 6 | 4 | 20 |
| sub06 | 19 | male | 9 | 13 | 8 | 18 | 8 | 4 | 7 | 16 | 7 | 2 | 17 | 11 |
| sub07 | 20 | female | 5 | 7 | 18 | 4 | 11 | 15 | 6 | 8 | 16 | 4 | 11 | 15 |
| sub08 | 22 | female | 3 | 16 | 11 | 3 | 13 | 14 | 4 | 16 | 10 | 2 | 14 | 14 |
| sub10 | 26 | female | 0 | 14 | 16 | 3 | 11 | 16 | 0 | 15 | 15 | 0 | 14 | 16 |
| sub14 | 20 | female | 2 | 15 | 13 | 1 | 14 | 15 | 0 | 15 | 15 | 0 | 12 | 18 |
| sub15 | 25 | female | 4 | 15 | 11 | 3 | 15 | 12 | 2 | 12 | 16 | 2 | 14 | 14 |
| sub18 | 18 | female | 0 | 8 | 15 | 0 | 3 | 20 | 0 | 4 | 23 | 0 | 1 | 29 |
| sub19 | 28 | male | 5 | 14 | 11 | 5 | 15 | 10 | 3 | 13 | 14 | 0 | 4 | 26 |
| sub21 | 21 | female | 2 | 7 | 21 | 2 | 7 | 21 | 0 | 10 | 20 | 1 | 9 | 20 |
| sub22 | 25 | female | 4 | 20 | 6 | 3 | 18 | 9 | 4 | 21 | 5 | 2 | 18 | 10 |
| sub24 | 22 | female | 0 | 7 | 23 | 0 | 9 | 21 | 0 | 11 | 19 | 0 | 10 | 20 |
| sub25 | 30 | male | 0 | 14 | 16 | 0 | 10 | 20 | 2 | 8 | 20 | 2 | 10 | 18 |
| sub26 | 21 | female | 2 | 8 | 19 | 1 | 7 | 22 | 1 | 3 | 26 | 0 | 6 | 24 |
| sub27 | 27 | female | 8 | 10 | 12 | 9 | 7 | 14 | 5 | 4 | 21 | 2 | 9 | 19 |
| sub28 | 24 | female | 3 | 20 | 7 | 5 | 17 | 8 | 6 | 11 | 13 | 2 | 18 | 10 |
| sub29 | 22 | female | 2 | 11 | 17 | 5 | 5 | 20 | 3 | 6 | 21 | 0 | 7 | 23 |
| sub30 | 28 | male | 6 | 14 | 10 | 7 | 5 | 18 | 3 | 17 | 10 | 2 | 18 | 10 |
| sub31 | 24 | female | 2 | 16 | 12 | 1 | 17 | 12 | 0 | 15 | 15 | 0 | 12 | 18 |

| Subject ID | Age (years) | Gender | Self-generated, Disrupted feedback | | | Self-generated, Fluent feedback | | | Externally generated, Disrupted feedback | | | Externally generated, Fluent feedback | | |
|---|---|---|---|---|---|---|---|---|---|---|---|---|---|---|
| | | | **Remember** | **Know** | **Forgotten** | **Remember** | **Know** | **Forgotten** | **Remember** | **Know** | **Forgotten** | **Remember** | **Know** | **Forgotten** |
| sub32 | 21 | female | 14 | 11 | 5 | 7 | 14 | 9 | 9 | 10 | 11 | 15 | 10 | 5 |
| sub33 | 31 | female | 5 | 15 | 10 | 8 | 14 | 8 | 1 | 19 | 10 | 0 | 13 | 17 |
| sub34 | 20 | male | 3 | 15 | 12 | 7 | 13 | 10 | 2 | 8 | 20 | 2 | 6 | 22 |
| sub35 | 24 | female | 4 | 13 | 13 | 2 | 11 | 17 | 1 | 14 | 15 | 0 | 14 | 16 |
| sub36 | 26 | female | 10 | 8 | 12 | 10 | 7 | 13 | 8 | 7 | 15 | 4 | 6 | 20 |
| sub37 | 18 | female | 1 | 12 | 17 | 1 | 14 | 15 | 2 | 11 | 17 | 0 | 6 | 24 |
| sub38 | 20 | female | 1 | 24 | 5 | 2 | 25 | 3 | 1 | 18 | 11 | 0 | 14 | 16 |
| sub41 | 19 | female | 5 | 15 | 10 | 4 | 17 | 9 | 2 | 16 | 12 | 2 | 11 | 17 |
| sub42 | 19 | female | 1 | 3 | 26 | 1 | 9 | 20 | 0 | 5 | 25 | 0 | 0 | 30 |
| sub43 | 18 | female | 1 | 14 | 15 | 0 | 12 | 18 | 1 | 10 | 19 | 1 | 14 | 15 |
| sub44 | 18 | female | 7 | 17 | 6 | 3 | 16 | 11 | 3 | 18 | 9 | 0 | 16 | 14 |
| sub45 | 18 | female | 1 | 9 | 20 | 2 | 12 | 16 | 0 | 7 | 23 | 2 | 3 | 25 |
| sub46 | 19 | male | 2 | 12 | 16 | 1 | 13 | 16 | 2 | 11 | 17 | 0 | 12 | 18 |
| sub47 | 18 | female | 2 | 14 | 14 | 1 | 10 | 19 | 0 | 9 | 21 | 0 | 9 | 21 |
| sub48 | 19 | male | 8 | 13 | 9 | 9 | 13 | 8 | 10 | 11 | 9 | 7 | 12 | 11 |
| sub49 | 22 | male | 7 | 9 | 14 | 2 | 10 | 18 | 2 | 9 | 19 | 1 | 6 | 23 |
| sub50 | 23 | female | 4 | 18 | 8 | 4 | 14 | 12 | 1 | 14 | 15 | 1 | 12 | 17 |
| sub51 | 22 | female | 3 | 13 | 14 | 1 | 9 | 20 | 1 | 13 | 16 | 0 | 9 | 21 |
| sub53 | 21 | female | 14 | 12 | 4 | 14 | 12 | 4 | 7 | 13 | 10 | 8 | 10 | 12 |
| sub54 | 22 | female | 0 | 19 | 11 | 2 | 19 | 9 | 3 | 9 | 18 | 0 | 13 | 17 |
| sub55 | 24 | male | 3 | 17 | 10 | 9 | 17 | 4 | 4 | 13 | 13 | 1 | 17 | 12 |

| Subject ID | Age (years) | Gender | Self-generated, Disrupted feedback | | | Self-generated, Fluent feedback | | | Externally generated, Disrupted feedback | | | Externally generated, Fluent feedback | | |
|---|---|---|---|---|---|---|---|---|---|---|---|---|---|---|
| | | | Remember | Know | Forgotten | Remember | Know | Forgotten | Remember | Know | Forgotten | Remember | Know | Forgotten |
| sub56 | 24 | male | 13 | 12 | 5 | 14 | 8 | 8 | 9 | 11 | 10 | 5 | 14 | 11 |
| sub57 | 24 | male | 0 | 18 | 12 | 4 | 18 | 8 | 1 | 11 | 18 | 1 | 7 | 22 |
| **Mean (SD)** | **22.21 (3.41)** | **31 F; 11 M** | **4.07 (3.74)** | **13.12 (4.23)** | **12.62 (5.05)** | **4.36 (4.17)** | **12.19 (4.42)** | **13.29 (5.36)** | **2.81 (2.80)** | **11.55 (4.26)** | **15.57 (4.95)** | **1.83 (2.88)** | **10.52 (4.61)** | **17.64 (5.46)** |

Counts denote retained action-onset-locked EEG epochs from the 42 participants included in the formal analyses. The final row reports the mean (SD) number of retained epochs across participants. Zero counts were retained when a participant contributed no trials to a recognition-response category. The sample included 31 female and 11 male participants. “Not Presented” trials were excluded.

**Table S2. Retained action-onset-locked EEG trial counts by experimental condition in Experiment 3**

*Related to Methods.*

| Subject ID | Age (years) | Gender | Self-generated, Disrupted feedback | Self-generated, Fluent feedback | Externally generated, Disrupted feedback | Externally generated, Fluent feedback |
|---|---|---|---|---|---|---|
| sub05 | 21 | female | 40 | 40 | 40 | 40 |
| sub06 | 19 | male | 40 | 40 | 40 | 40 |
| sub07 | 20 | female | 40 | 40 | 40 | 40 |
| sub08 | 22 | female | 40 | 40 | 40 | 40 |
| sub10 | 26 | female | 40 | 40 | 40 | 40 |
| sub14 | 20 | female | 40 | 40 | 40 | 40 |
| sub15 | 25 | female | 40 | 40 | 40 | 40 |
| sub18 | 18 | female | 29 | 26 | 30 | 35 |
| sub19 | 28 | male | 40 | 40 | 40 | 40 |
| sub21 | 21 | female | 40 | 40 | 40 | 40 |
| sub22 | 25 | female | 40 | 40 | 40 | 40 |
| sub24 | 22 | female | 40 | 40 | 40 | 40 |
| sub25 | 30 | male | 40 | 40 | 40 | 40 |
| sub26 | 21 | female | 39 | 40 | 40 | 40 |
| sub27 | 27 | female | 40 | 40 | 40 | 40 |
| sub28 | 24 | female | 40 | 40 | 40 | 40 |
| sub29 | 22 | female | 40 | 40 | 40 | 40 |
| sub30 | 28 | male | 40 | 40 | 40 | 40 |
| sub31 | 24 | female | 40 | 40 | 40 | 40 |
| sub32 | 21 | female | 40 | 40 | 40 | 40 |
| sub33 | 31 | female | 40 | 40 | 40 | 40 |
| sub34 | 20 | male | 40 | 40 | 40 | 40 |
| sub35 | 24 | female | 40 | 40 | 40 | 40 |
| sub36 | 26 | female | 40 | 40 | 40 | 40 |
| sub37 | 18 | female | 40 | 40 | 40 | 40 |
| sub38 | 20 | female | 40 | 40 | 40 | 40 |
| sub41 | 19 | female | 40 | 40 | 40 | 40 |
| sub42 | 19 | female | 40 | 40 | 40 | 40 |
| sub43 | 18 | female | 40 | 40 | 40 | 40 |
| sub44 | 18 | female | 40 | 40 | 40 | 40 |
| sub45 | 18 | female | 40 | 40 | 40 | 40 |
| sub46 | 19 | male | 40 | 40 | 40 | 40 |
| sub47 | 18 | female | 40 | 40 | 40 | 40 |
| sub48 | 19 | male | 40 | 40 | 40 | 40 |

| Subject ID | Age (years) | Gender | Self-generated, Disrupted feedback | Self-generated, Fluent feedback | Externally generated, Disrupted feedback | Externally generated, Fluent feedback |
|---|---|---|---|---|---|---|
| sub49 | 22 | male | 40 | 40 | 40 | 40 |
| sub50 | 23 | female | 40 | 40 | 40 | 40 |
| sub51 | 22 | female | 40 | 40 | 40 | 40 |
| sub53 | 21 | female | 40 | 40 | 40 | 40 |
| sub54 | 22 | female | 40 | 40 | 40 | 40 |
| sub55 | 24 | male | 40 | 40 | 40 | 40 |
| sub56 | 24 | male | 40 | 40 | 40 | 40 |
| sub57 | 24 | male | 40 | 40 | 40 | 40 |
| **Mean (SD)** | **22.21 (3.41)** | **31 F; 11 M** | **39.71 (1.70)** | **39.67 (2.16)** | **39.76 (1.54)** | **39.88 (.77)** |

Counts denote retained action-onset-locked EEG epochs from the 42 participants included in the formal analyses. The final row reports the mean (SD) number of retained epochs across participants. The sample included 31 female and 11 male participants. Marker prefixes 11, 12, 21, and 22 corresponded to self-generated–disrupted feedback, self-generated–fluent feedback, externally generated–disrupted feedback, and externally generated–fluent feedback, respectively.
